# Elucidating DNA-binding protein dynamics in *Salmonella* Typhimurium within macrophages using a breakthrough low-input ChIP-exo approach

**DOI:** 10.1101/2024.06.20.599816

**Authors:** Joon Young Park, Minchang Jang, Eunna Choi, Sang-Mok Lee, Ina Bang, Jihoon Woo, Seoyeon Kim, Eun-Jin Lee, Donghyuk Kim

**Affiliations:** School of Energy and Chemical Engineering, Ulsan National Institute of Science and Technology (UNIST), Ulsan, 44919, Republic of Korea; Department of Life Sciences, College of Life Sciences and Biotechnology, Korea University, Seoul, 02841, Republic of Korea

**Author notes:** These authors contributed equally to this work (Joon Young Park, Minchang Jang). Correspondence should be addressed to D.K. and E.-J.L.

## Abstract

Genome-wide identification of binding profiles for DNA-binding proteins in *Salmonella* Typhimurium within macrophages is crucial for understanding virulence gene expression and cellular processes. However, this task remains challenging due to the limited amount of intracellular bacterial cells. Here, we present ChIP-mini, a low-input ChIP-exo utilizing 5,000-fold reduced number of initial bacterial cells and analysis pipeline (DiffExo), to identify genome-wide binding dynamics of DNA-binding proteins in host-infected pathogens. Applying ChIP-mini to intracellular *S.* Typhimurium, we identified 642 and 1,837 binding sites of H-NS and RpoD, respectively, with near-base pair resolution, elucidating their roles in transcriptional initiation. Post-infection, we observed 21 significant reductions in H-NS binding at intergenic regions, exposing the promoter region of virulence genes, such that those in *Salmonella* pathogenicity islands-2, 3 and effectors, facilitating RpoD binding for transcription initiation. Furthermore, we identified 24 novel RpoD binding sites and 19 significantly increased RpoD bindings at the transcription start sites of virulence genes. These findings substantially enhance our understanding of how H-NS and RpoD simultaneously coordinate the transcription initiation of virulence genes within macrophages. Collectively, this optimized method demonstrates a tool that can be broadly adapted to elucidate transcriptional regulatory networks of host-infected pathogens, revealing critical interactions between host and microbe.

## Introduction

*Salmonella enterica* serovar Typhimurium is a nontyphoidal Gram-negative pathogen responsible for worldwide foodborne illnesses, primarily causing gastroenteritis, and is capable of infecting a wide range of host cell types ^1^. Notably, macrophages, which serve as the primary defense against invading bacteria, are also crucial colonization niches of *S.* Typhimurium ^2^. The ability of pathogen to survive and replicate within macrophages facilitates its establishment of systemic disease in susceptible hosts ^2-4^. Understanding this ability requires elucidating the transcriptional regulatory networks (TRNs) of *S.* Typhimurium within macrophages, a complex challenge that involves understanding host-microbe interactions from a transcriptional regulation standpoint. Revealing these complex networks is essential for interpreting virulence mechanisms of pathogens in response to environmental stimuli.

Recent studies have investigated the transcript expression profiling of *S.* Typhimurium under macrophage intracellular and infection-relevant *in vitro* growth conditions ^5-7^. Additionally, RpoE binding profiling of *S.* Typhimurium within macrophages was reported using chromatin immunoprecipitation-seq (ChIP-seq) with limited resolution ^8^. Despite these efforts, obtaining high-resolution binding profiles to investigate the genome-wide binding dynamics of DNA-binding proteins within macrophages remains elusive due to the frequently limited number of S. Typhimurium recovered from infection studies.

To gain insight into the transcriptional regulatory mechanisms of *S.* Typhimurium virulence genes during infection, two DNA-binding proteins, H-NS and RpoD, were selected to elucidate their binding dynamics. H-NS, also known as histone-like nucleoid structuring protein, acts as a global transcription silencer in Gram-negative bacteria ^9-11^. H-NS binds to AT-rich DNA associated with horizontally-acquired genes, including *Salmonella* pathogenicity islands (SPIs) and various virulence genes, playing a pivotal role in alleviating fitness costs by suppressing those genes under normal conditions ^9,12^. Additionally, RpoD, the housekeeping sigma factor, has been extensively studied for recognizes the majority of promoters in *S.* Typhimurium and *Escherichia coli* in previous research ^13-17^. The predominant binding sites of RpoD suggest its significant role in governing the transcription initiation of various virulence genes. However, despite their importance, detailed genome-wide binding maps and binding dynamics of these DNA-binding proteins are still largely unknown within host cells.

Over the past two decades, ChIP methodology has been combined with whole-genome DNA microarray (ChIP-chip) or deep sequencing (ChIP-seq) to elucidate the genome-wide binding profiles of DNA-binding proteins ^18,19^. However, these methods suffer from low resolution due to the size heterogeneity of randomly sheared immunoprecipitated-DNA (IP-DNA) required for alignment to the genome, hindering the identification of complete and high-resolution binding sites ^20^. To address this limitation, ChIP-exo (Chromatin immunoprecipitation with exonuclease treatment) was developed. This technique uses lambda exonuclease to digest the 5’ end of IP-DNA up to the protein-DNA crosslinking point, achieving near-base pair resolution ^20,21^.

ChIP-exo has been further modified for bacterial use, enabling the identification of *in vivo* binding profiles of transcription factors (TFs) on the bacterial genome ^15,16,22-24^. However, a significant drawback of the current ChIP-exo method is its requirement for a large quantity of bacterial cells (in the 10^10^∼10^11^ range) due to the complex processes and the short IP-DNA, which limits its application to host intracellular bacteria. Therefore, an optimized method was necessary that could utilize a low number of initial bacterial cells for studying the TRNs of *S.* Typhimurium within macrophages.

Here, we present the ChIP-mini, a method suitable for working with as few as 4.8×10^6^ bacterial cells, which faithfully maintain relevant binding profiles generated by the traditional ChIP-exo method. This method was successfully applied during the traditional infection process of macrophages with *S.* Typhimurium to capture both host extracellular and intracellular bacteria (*in vitro* and *in vivo* ChIP-mini). We demonstrate the applications of ChIP-mini in studying the TRNs of *S.* Typhimurium within macrophages, achieving near-base pair resolution to detect the impact of environmental changes on the binding profiles of H-NS and RpoD.

For further validation, we constructed the DiffExo pipeline (ChIP-exo peak normalization pipeline) to identify binding profiles with statistically different intensities between macrophages extracellular and intracellular conditions. Through these efforts, we successfully discovered genome-wide binding maps of H-NS and RpoD with near-base pair resolution within macrophages. Utilizing the DiffExo pipeline, we identified alterations in the binding intensity of both DNA-binding proteins when *S.* Typhimurium transitions from macrophage extracellular to intracellular environments, offering insights into potential relationships for transcription initiation between H-NS and RpoD binding upstream of various virulence genes.

Collectively, this study marks a cornerstone toward elucidation of the TRNs of this infectious host intracellular pathogen and provides a valuable resource for future studies aiming at decoding the comprehensive genome-wide *in vivo* regulatory roles of various DNA-binding proteins.

## Results

### Optimized ChIP-mini method generates reproducible RpoD binding profiles using low-input bacterial cells

To ensure compatibility with a low number of initial bacterial cells, we optimized the procedures and reagent usage of the traditional ChIP-exo method (Figure S1A). ChIP-mini was modified in three major aspects compared to the traditional method: sonication for DNA fragmentation, the amount of reagents usage, and the IP-DNA purification method. Details of these modifications of ChIP-mini are presented in Supplementary Text S1.

To assess the reproducibility of ChIP-mini method, we determined RpoD binding profiles of *E. coli* K-12 MG1655 starting from different numbers of cells. Using the ChIP-mini method, sequencing libraries were constructed for *E. coli* RpoD binding profiling from six samples with progressively reduced numbers of initial bacterial cells. Notably, all libraries showed comparable amplification cycles to those of traditional ChIP-exo, indicating that ChIP-mini effectively minimized the loss of IP-DNA and maintained amplification cycles consistent with traditional ChIP-exo, even with a low number of initial bacterial cells (Figure 1C). Furthermore, these ChIP-mini libraries were verified for their consistent fragment distribution (Figure S1D), with no abnormalities observed compared to the traditional ChIP-exo library.

**Figure 1.**
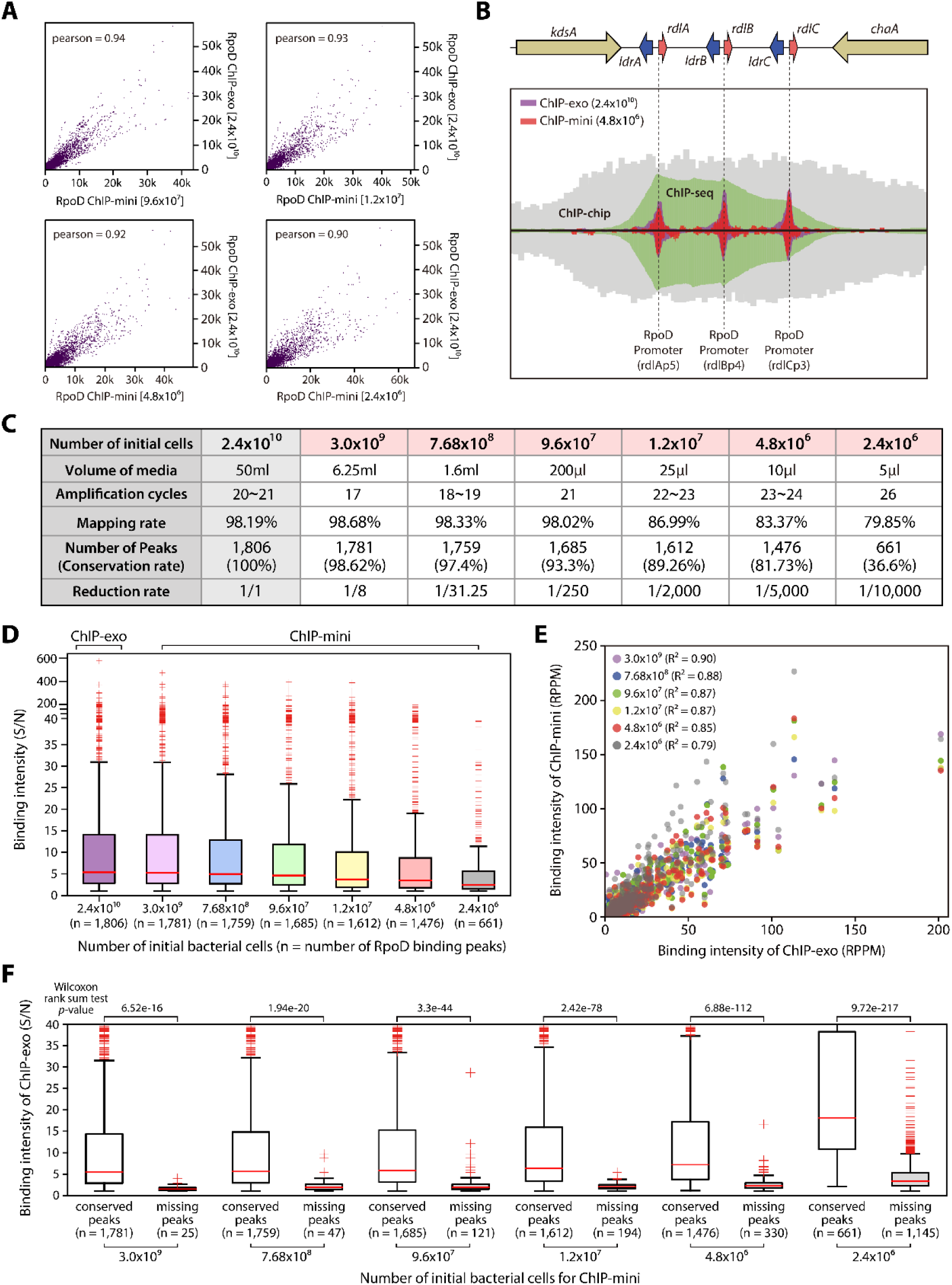
Comparative analysis of traditional ChIP-exo and enhanced ChIP-mini, preserving the advantages of ChIP-exo with a minimum input. (A) Scatter plot depicting the comparative analysis of the sequence alignment BAM files from traditional ChIP-exo and four ChIP-mini results. The read counts from each library were segmented into 10-bp bins across the *E. coli* K-12 MG1655 genome, with each dot in the plot symbolizing a specific genomic region. Abbreviation: Pearson’s correlation coefficient: pearson (B) Evaluation ChIP-mini against three other ChIP methods for delineating transcription factor binding profiles. ChIP-mini achieves near single base-pair resolution, even with a minimal initial quantity of bacterial cells (4.8×10^6^). (C) Assessment of mapping and conservation efficacy in ChIP-mini libraries in relation to decrease in initial bacterial cell count. (D) A boxplot illustrating the binding intensity of RpoD (signal-to-noise ratio, S/N) across ChIP-mini and traditional ChIP-exo datasets. (E) A scatter plot displaying the normalized binding intensity (RPPM) of the overlapping peaks identified in both traditional ChIP-exo and each ChIP-mini dataset. (F) Analysis of the binding intensities of both conserved and missing peaks. In the RpoD ChIP-exo dataset, peaks missing in ChIP-mini exhibited weaker intensities (rank-sum test *p*-value < 0.05).

Amplified libraries were sequenced to a depth of 8.40 ± 1.88 million reads per ChIP-mini sample, and the nucleotide frequency at the 5’ end of the sequencing tags was compared with traditional ChIP-exo library. Unlike ChIP-seq, ChIP-exo exhibit a characteristic found in paired-end read sequencing data. While ChIP-seq has only 5’ end resulting from adapter ligation after sonication in Read_1 and Read_2 file, ChIP-exo features an exonuclease-digested 5’ end in the Read_1 file (Figure S2A). A comparison of ChIP-exo with ChIP-mini data showed that ChIP-mini effectively maintained this ChIP-exo trait (Figure S2B). Additionally, Pearson correlation coefficients of aligned sequencing files (10 bp-bins) between traditional ChIP-exo and each ChIP-mini consistently exceeded 0.9, even in cases with less than 9.6 x 10^7^ datasets (Figure 1A) ^25^. A comparison of ChIP-exo, ChIP-mini, and other ChIP methods demonstrated that the ChIP-mini achieves near-base pair resolution for RpoD binding sites, which is an important advantage of ChIP-exo ^26^ (Figure 1B). Using a deep-learning based peak calling pipeline (DEOCSU), RpoD binding peaks were identified for traditional ChIP-exo, and six ChIP-mini libraries ^27^. While the mapping rate remained relatively consistent across all ChIP-mini cases, the peak conservation rate exhibited a significant decline in the 2.4×10^6^ dataset (Figure 1C). Therefore, we established 4.8×10^6^ as the threshold for the initial bacterial cell numbers in the ChIP-mini method.

PCR amplification from limited IP-DNA has been known to result in a reduction of mappable reads and the generation of undesirable reads, such as background-mapped and unmapped reads ^28^. To assess this issue, the binding intensity of RpoD binding peaks for each ChIP-mini dataset was calculated using signal-to-noise ratio (S/N). As predicted, the S/N value showed a gradual decrease in the ChIP-mini datasets (Figure 1D), with a notable decline observed in datasets below the threshold of 4.8×10^6^. This decline was attributed to an increased noise level relative to the total number of reads (Figure S2C). To validate the pattern of RpoD binding intensities between ChIP-exo and each ChIP-mini dataset, the binding intensity of overlapping binding peaks was normalized using RPPM (Reads Per Peak per Million). Interestingly, Pearson correlation coefficients of these normalized values revealed a strong correlation in the trend of binding intensities between each ChIP-mini dataset and the traditional ChIP-exo dataset (R^2^ > 0.79) (Figure 1E).

Moreover, to determine whether the reduction in the initial bacterial cell counts in ChIP-mini affects the major peaks with strong intensity, the binding intensity of missing peaks in each ChIP-mini dataset was calculated with traditional ChIP-exo dataset. Contrary to the conserved peaks, missing peaks from each ChIP-mini dataset exhibited significantly weaker binding intensities in ChIP-exo data (Figure 1F, rank-sum test *p*-value < 0.05), suggesting ChIP-mini can reproduce the majority of binding profiles of the traditional ChIP-exo method. Additionally, the RpoD binding site sequence motifs of ChIP-exo and ChIP-mini datasets were identified as ttgnca-15bp-gntAtaaT (Figure S2D, lower-case characters indicate an information content <1 bit). These motifs were identical to those found in previous studies ^13,15,29^. Thus, these results suggest that ChIP-mini has strong feasibility for generating genome-wide binding profiles using a significantly smaller number of bacterial cells, up to 5,000-fold less than traditional ChIP-exo. Therefore, we confirmed the compatibility of ChIP-mini in limited environmental conditions, such as host infection, aiming to identify high-resolution binding profiles of DNA-binding proteins from both host extracellular and intracellular bacteria.

### Applications of ChIP-mini for *in vivo S.* Typhimurium infection inside macrophages

To determine whether ChIP-mini can be identify changes in the binding patterns of bacterial DNA-binding regulatory proteins during host infection, macrophage-like cell line (J774A.1) and *S.* Typhimurium were employed in the infection model. Additionally, in order to statistically discern changes in binding intensity during host cell invasion, we constructed a ChIP-mini analysis workflow incorporating the DiffExo pipeline (Figure 2A). The DiffExo pipeline was designed to perform the following tasks: 1) identification of differentially binding sites between two or more sample groups using DEseq2, 2) normalization of binding intensity, 3) statistics analysis, and 4) generation of plots (Figure S3) ^30,31^.

**Figure 2.**
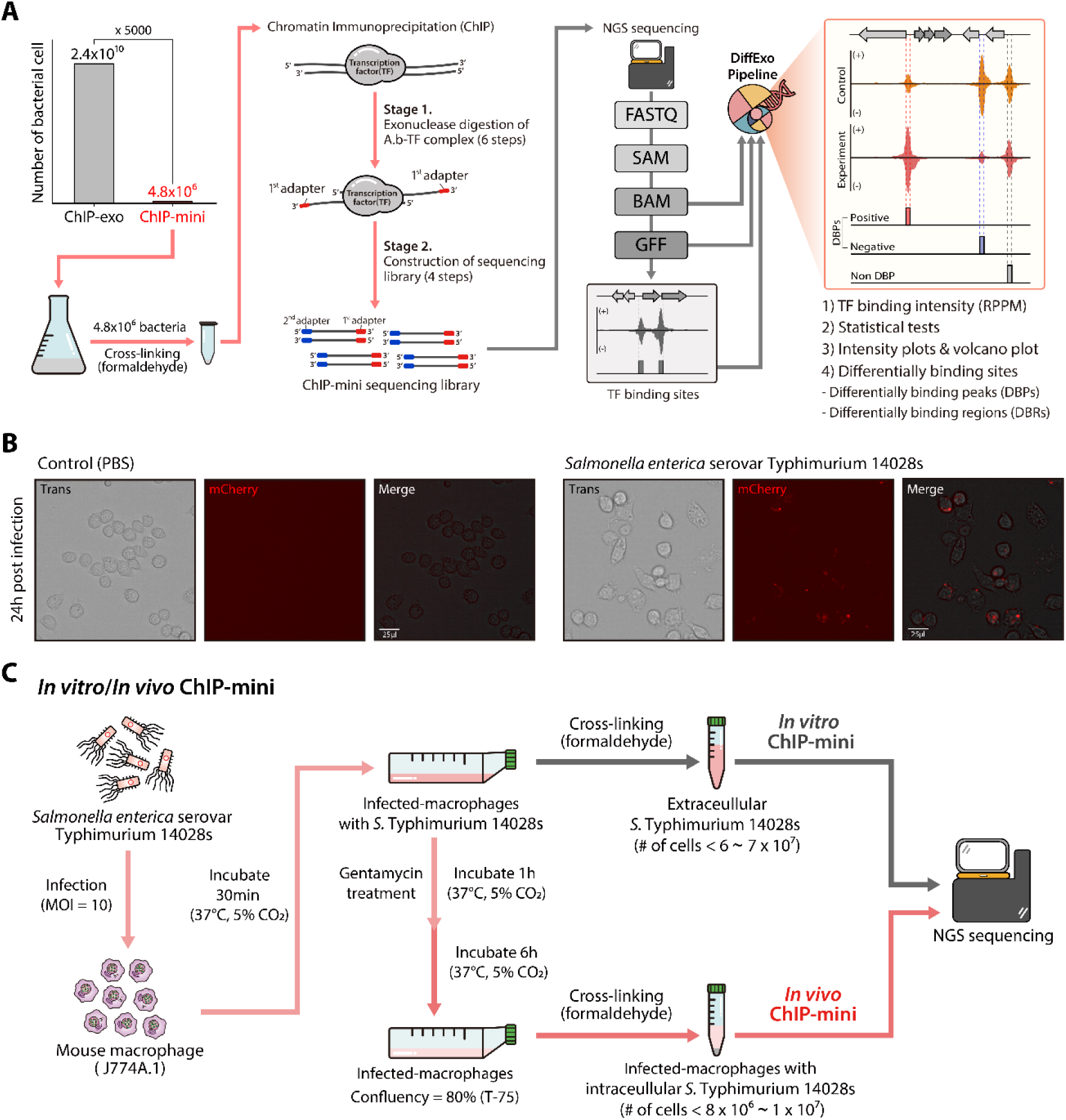
Optimized ChIP-mini method to identify binding sites of DNA-binding proteins in *S.* Typhimurium 14028s under macrophage extracellular and intracellular conditions. (A) Schematic diagram of the ChIP-mini workflow: this method has been optimized to use with a minimal quantity of initial bacterial cells. The DiffExo analysis pipeline was utilized for normalizing binding intensity and identifying differential binding sites between the control and the experimental ChIP-mini datasets. (B) Macrophage-like cells (J774A.1) underwent infection with *S.* Typhimurium carrying pFCcGi, which constitutively expresses mCherry. Phosphate buffered saline (PBS) was used as a control. At 24hr post-infection (hpi) the infected samples were fixed, and the presence of *S.* Typhimurium were visualized through the mCherry fluorescence (red). (C) Schematic diagram illustrating the *in vitro* and *in vivo* ChIP-mini experiments with macrophage extracellular and intracellular *S.* Typhimurium.

To assess infection efficiency and estimate the number of intracellular bacteria, *S.* Typhimurium harboring the pFCcGi vector, which constitutively expresses mCherry was used (Figure 2B). Following the infection of this strain into the macrophages, *S.* Typhimurium demonstrated an approximate 90% infection efficiency. This suggests that nearly every macrophage contained one or more intracellular *S.* Typhimurium (CFU > 8×10^6^). This high infection efficiency indicates that the number of intracellular *S.* Typhimurium can exceed the ChIP-mini threshold (4.8×10^6^), allowing for the harvesting of a sufficient number of bacterial cells to perform ChIP-mini during macrophage infection experiments.

Previous studies have demonstrated that while H-NS amounts remain constant under laboratory growth conditions ^32^, while *S.* Typhimurium decreases H-NS amounts via Lon and PhoP to activate the transcription of numerous virulence-related HTGs during infection, facilitating successful survival within macrophages ^9,33^. However, genome-wide changes in H-NS binding coupled with RpoD binding profiles for elucidating transcriptional initiation mechanisms inside macrophages are still unknown. Therefore, to identify genome-wide dynamic changes in H-NS and RpoD binding profiles during macrophage invasion, we designed two types of ChIP-mini applications: *in vitro* ChIP-mini and *in vivo* ChIP-mini (Figure 2C). For these applications, *S.* Typhimurium *hns*-8myc tagged strain for H-NS and the wild-type strain for RpoD were used to infect macrophages, respectively. After infection, extracellular bacteria in DMEM were crosslinked and isolated to identify macrophage extracellular H-NS and RpoD binding profiles (*in vitro* ChIP-mini). At 6 hours post-infection (hpi), the *S.* Typhimurium-infected macrophages were co-crosslinked. The co-crosslinked intracellular bacteria were then utilized to identify macrophage intracellular H-NS and RpoD binding profiles (*in vivo* ChIP-mini).

Following both ChIP-mini experiments, libraries for H-NS and RpoD under both conditions were constructed. Sequentially, these libraries were aligned to *S.* Typhimurium genome and the host genome (*Mus musculus*) to check for host nucleic acid contamination (Figure S4A). In the *in vitro* libraries, the majority of sequencing reads aligned to the *S.* Typhimurium genome. However, in the *in vivo* libraries, host DNA contamination due to the co-crosslinking method had a more pronounced impact, resulting in a lower mapping rate compared to the *in vitro* libraries. Despite this lower mapping rate, approximately 2 million reads were obtained for the *in vivo* RpoD libraries, which is sufficient for subsequent analysis, as demonstrated in previous bacterial studies ^22,23,34^. To investigate whether a lower number of initial *S.* Typhimurium leads to lower library complexity, we assessed library complexity by calculating the number of uniquely mapped reads (Figure S4B). Uniquely mapped reads represented over 87% across all aligned libraries, suggesting that high library complexity is maintained even with a limited number of initial bacterial cells. Additionally, Pearson correlation coefficients of H-NS and RpoD aligned sequencing files (10 bp-bins) between technical replicates were consistently above 0.99 in all libraries, indicating a high level of reproducibility (Figure S4C). Thus, the ChIP-mini applications successfully produced eight sequencing libraries for identifying genome-wide binding profiles of H-NS and RpoD in S. Typhimurium under both macrophage extracellular and intracellular conditions.

### Genome-wide binding profiles of H-NS under macrophage extracellular and intracellular conditions

Previously, several H-NS binding regions in *Salmonella* have been characterized through *in vitro* DNA-binding experiments and ChIP methods, such as ChIP-chip and ChIP-seq ^9,35-38^. However, despite the importance of H-NS in regulating the transcription of virulence genes, detailed information about the change in H-NS binding patterns and intensity during macrophage infection remains elusive. Additionally, the low sensitivity due to the broad binding regions of H-NS oligomers further complicates the identification of precise fluctuating regions in response to environmental shifts. To address these challenges, we utilized ChIP-mini applications, which offer improved resolution in dissecting binding regions compared to ChIP-seq (Figure S5A). This allowed us to determine genome-wide binding profiles of H-NS in *S.* Typhimurium under macrophage extracellular and intracellular conditions with high resolution. From the genome-wide binding profile of H-NS ChIP-mini datasets, a total of 642 H-NS binding regions were identified under both conditions (Figure 3A and Supplementary Table S1). Consistent with previous research, these binding regions occupied 16.6% of the genome in terms of base pairs, exhibiting an average GC content of 43.7%, which is lower than that of the entire *S.* Typhimurium 14028s genome (52.1%) ^9,39^.

**Figure 3.**
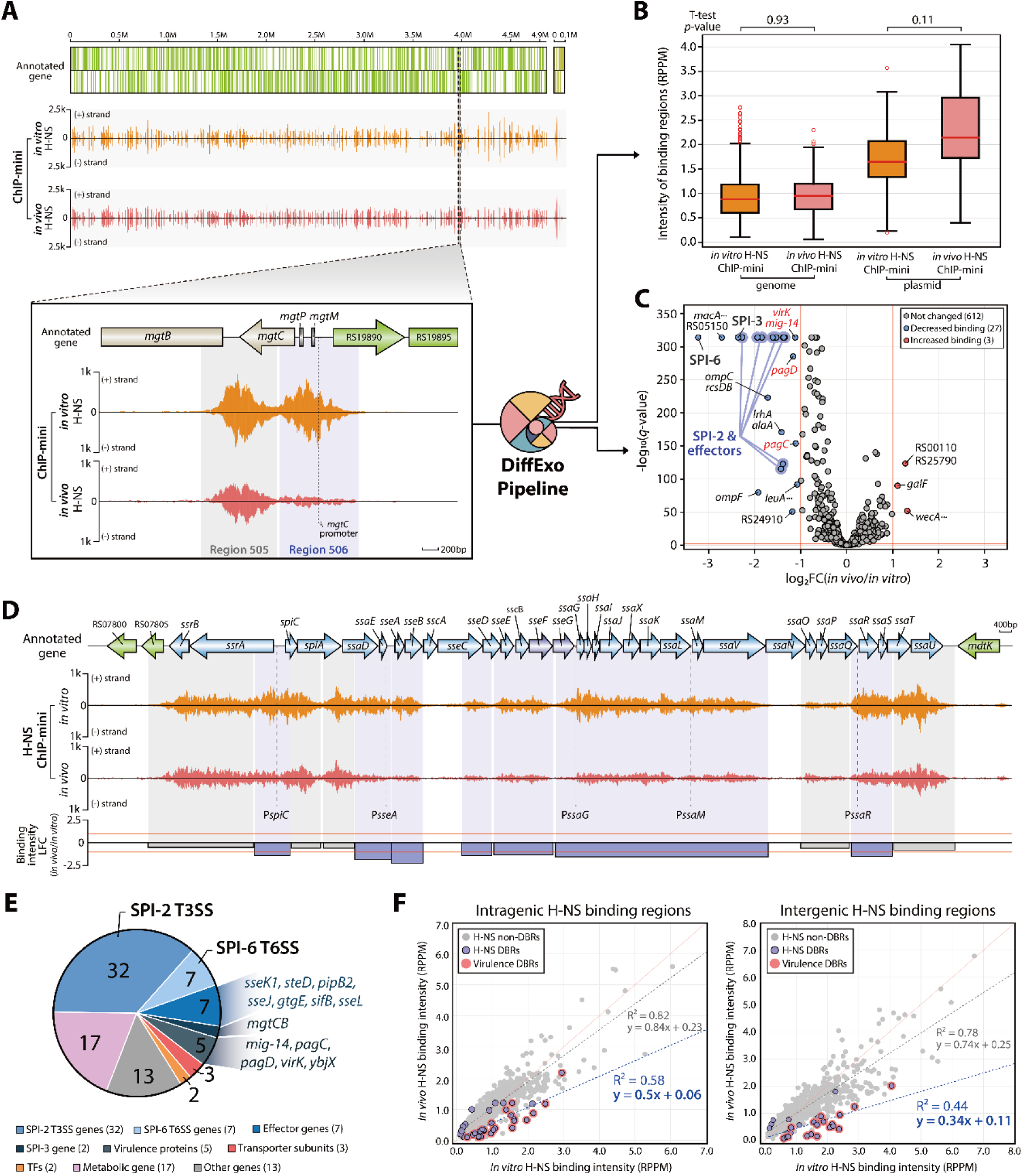
Genome-wide identification of changes in H-NS binding intensity when *S.* Typhimurium transitioning from macrophage extracellular to intracellular conditions. (A) A comprehensive landscape of H-NS binding patterns throughout the *S.* Typhimurium 14028s genome and its plasmid under macrophage extracellular to intracellular conditions. A detailed inset emphasizes a specific example of variations in H-NS binding intensity due to environmental changes. (B) Normalized binding intensities for H-NS binding regions under extracellular and intracellular conditions were calculated (t-test *p*-value > 0.05). (C) Volcano plot displaying H-NS differentially bound regions (DBRs) as response to environmental changes (log_2_ fold change ≤ -1 or ≥ 1, and false discovery rate < 0.05). Blue lines indicate DBRs associated with SPI-2 and effector genes, while red font emphasizes genes related to virulence. (D) Variations in H-NS binding intensity within the SPI-2 region of *S.* Typhimurium. Red lines represent the threshold for DBRs, marked at -1 and 1 in binding intensity LFC. Blue boxes highlight negative DBRs, and grey boxes represent areas with no significant changes in binding (non-DBRs). In the ChIP-mini data, ‘(+)’ and ‘(−)’ indicate reads mapped onto forward and reverse strands, respectively. (E) A pie chart illustrating the proportion of target genes linked to both positive and negative H-NS DBRs. (F) The H-NS binding regions were categorized into intragenic and intergenic regions to assess the extent of impact by macrophages intracellular conditions. Pearson correlation coefficients and trend lines of H-NS binding intensities were derived from the comparative analysis of *in vitro* and *in vivo* ChIP-mini datasets.

To gain further insight into the target genes associated with H-NS, the integration of H-NS binding region information with transcription units (TUs) annotation became necessary. Thus, the TUs with H-NS binding sites in their upstream regulatory region were chosen from the reported TU annotation ^13,40^. In addition, intragenic H-NS binding region was considered to influence only that gene in which they are located. Currently, a total of 745 genes have been characterized with strong evidence as H-NS-associated genes by ChIP-chip ^9^. Using our ChIP-mini datasets, we significantly expanded the number of H-NS-associated genes to comprise 1,242 genes in 787 TUs (Supplementary Table S1). Moreover, H-NS bindings were identified in six *Salmonella* pathogenicity islands (SPIs). In SPI-1 and SPI-2, the most representative SPIs, H-NS binding regions were divided into 16 and 12, respectively (Figure S6). Additionally, a total of six, nine, two, and eight binding regions were identified in SPI-3, SPI-4, SPI-5, and SPI-6, respectively (Figure S7). Furthermore, H-NS was found to bind to 32 genes associated with the type III secretion system (T3SS:29) and type VI secretion system (T6SS:3) effectors (Figure S8).

To facilitate the comparison of H-NS binding intensity between macrophages extracellular and intracellular conditions, it was imperative to confirm the homogeneity of the ChIP-mini datasets. Thus, the trend of noise level relative to the total number of reads was calculated using traditional ChIP-exo and both ChIP-mini datasets, revealing strong correlations across all datasets (R^2^ > 0.98) (Figure S5B). This robust correlation implies that H-NS ChIP-mini datasets were uniformly generated under both conditions, enabling reliable comparative analysis. Therefore, normalized binding intensities were calculated using the DiffExo pipeline. The overall distribution of H-NS binding intensities from both conditions was similar in both the genome and plasmid (T-test *p*-value > 0.05) (Figure 3B). Additionally, Pearson correlation coefficients of normalized binding intensity between both conditions showed high correlation (R^2^ > 0.8), indicating substantial similarity in binding intensity between the two conditions (Figure S5C). Consequently, these results demonstrate that ChIP-mini applications generated reproducible genome-wide H-NS binding regions of *S.* Typhimurium under both conditions. In addition, the high similarity between the *in vitro* and *in vivo* ChIP-mini datasets raises the question of whether there are regions where the intensity of H-NS binding significantly differs.

### H-NS binding intensity significantly decreases in SPI-2 and effector genes after infection

To determine the differentially binding regions (DBRs) of H-NS, we conducted a comparative analysis using the DiffExo pipeline. Overall, a total of 30 differentially binding regions were identified within macrophages (Supplementary Table S2, absolute value of log_2_ intensity fold change ≥ 1.0 and false discovery rate < 0.05). Of these, 27 regions were classified as negative DBRs, demonstrating a significant decrease in binding intensity, while three DBRs were classified as positive DBRs, exhibiting increased binding intensity. Additionally, the changes in binding intensities were more significant in the negative DBRs (T-test *p*-value < 0.05), indicating that decreases in binding intensity were more pronounced than increases within macrophage (Figure S5D). Furthermore, our analysis revealed that 21 out of 27 negative DBRs are associated with virulence genes, highlighting the importance in H-NS binding for the virulence of *S.* Typhimurium within macrophages (Figure 3C).

In *S.* Typhimurium, SPI-1 and SPI-2 are crucial for the invasion of eukaryotic cells and for facilitating replication to cause systemic disease within host cells, respectively ^41^. However, the T3SS proteins of SPI-1 are recognized by macrophages, triggering a caspase-1-mediated form of inflammatory cell death known as pyroptosis ^42^. To avoid inducing pyroptosis, late-stationary phase (LSP) cultures are commonly used in macrophage infection studies, during which *Salmonella* expresses SPI-1 genes at low levels ^43^. Therefore, the repression of SPI-1 is crucial under both macrophage extracellular and intracellular conditions, while SPI-2 activation is specifically essential for replication and proliferation of *S.* Typhimurium within macrophages ^41^. Notably, a significant reduction in H-NS binding intensity was particularly evident in SPI-2 but not SPI-1, and these negative DBRs included all transcription start sites (TSS) of SPI-2 that have been previously reported ^6^ (Figure 3D). Additionally, a single negative DBR was found in the intergenic region of SPI-3 and SPI-6 (Figure S7).

The target genes associated with these 30 DBRs encompassed 88 target genes, with about 60.2% identified as virulence-related genes (Figure 3E and Supplementary Table S3). These target genes included the entirety of SPI-2 genes, a portion of SPI-3 and SPI-6 genes. In SPI-3, a negative DBR in SPI-3 was found upstream of *mgtCB* transcriptional unit (Figure S7), genes that promote *Salmonella* survival within macrophages ^44,45^. Additionally, a negative DBR was located in the promoter region of *tagKJ*-*tssEFGA* and *tssH* transcriptional units. Moreover, seven effector genes were identified among the negative DBR target genes (Figure S9). Six of these effector genes have been reported to play a vital role in *Salmonella* virulence within host cells. SseK1 inhibits TNFα-induced NF-kB signaling of innate immune signaling ^46^. SseJ and PipB2 contribute to the formation and elongation of Salmonella-induced tubular (SITs), which are tubular extensions of the *Salmonella*-containing vacuole (SCV) ^47-50^. GtgE is a cysteine protease that prevents the accumulation of Rab29, Rab32, and Rab38 on SCV and SITs ^51,52^. In addition, SteD is a transmembrane effector required for the depletion of surface mature major histocompatibility class II (mMHCII), thereby inhibiting antigen presentation and T-cell proliferation within dendritic cells ^53^. Presumably, SseL functions as a deubiquitinase, preventing the accumulation of lipid droplets and inhibiting the clearance of cytosolic aggregates, leading to late macrophage cell death ^54-56^. The remaining effector, SifB, is a core effector always found in intestinal serovars, suggesting its significance in *Salmonella* virulence, although its precise function within macrophages remains elusive ^48^. Additionally, a substantial decrease in H-NS binding intensity was also detected in three other important virulence genes (*mig-14*, *pagC*, and *virK*) ^57-59^. Furthermore, to identify which genomic regions exhibited a significant decrease in binding intensity, we divided H-NS binding regions into intragenic and intergenic categories and compared the changes in binding intensity (Figure 3F). This analysis revealed that the reduction in H-NS binding intensity was more pronounced in intergenic regions. Altogether, these findings suggest that a reduction in H-NS binding at the TSS regions may increase the accessibility of sigma factors or other TFs, facilitating the transcription initiation of virulence genes under macrophage intracellular conditions.

### Dynamic changes in RpoD binding profiles in response to environmental shift

To ascertain whether the reduced binding of H-NS in response to environmental shift induces transcription initiation, we identified the binding sites of RpoD under both conditions using ChIP-mini applications. These ChIP-mini datasets showed a high correlation in the trend of noise level relative to the total number of reads across all ChIP-mini and traditional ChIP-exo datasets (R^2^ > 0.98), indicating the robustness of the data for further analysis (Figure S10A).

A total of 1,978 and 1,837 RpoD binding sites were identified under macrophage extracellular and intracellular conditions, respectively (Figure 4A and Supplementary Table S1). The majority of these binding sites overlapped, and macrophage intracellular RpoD binding sites in the genome exhibited higher intensity compared to extracellular conditions (Figure S10B, rank-sum test *p*-value < 0.05). Interestingly, the normalized binding intensities of overlapping peaks exhibited a low correlation, suggesting that RpoD binding at these peaks undergoes changes within macrophages (Figure S10C). Integrating TU information with RpoD binding sites in their upstream regulatory regions, we identified 1,404 TUs with 2,123 genes and 1,306 TUs with 2,005 genes in each ChIP-mini dataset (Figure 4B). The sequence motifs derived from RpoD binding sites were also found to be "ttgaca-15bp-gntAtaaT," consistent with traditional ChIP-exo of *E. coli* and previous finding (Figure 4C, lower-case characters indicate an information content <1 bit) ^16^.

**Figure 4.**
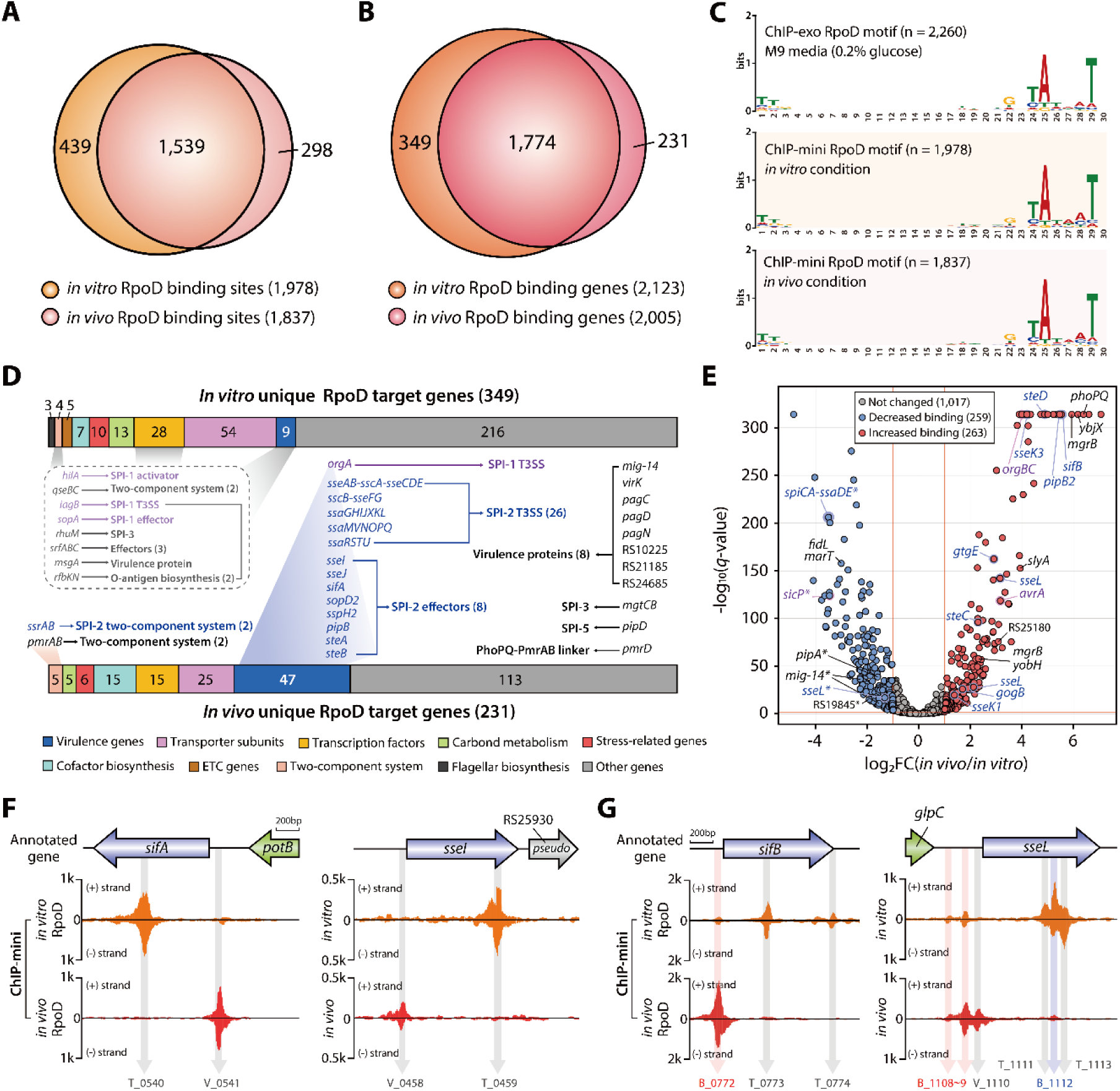
Genome-wide identification of changes in RpoD binding site and intensity during transition from macrophage extracellular to intracellular conditions. (A) Analysis of the overlaping RpoD binding sites between macrophages extracellular to intracellular conditions. (B) Comparison of RpoD target genes identified by ChIP-mini under macrophage conditions, both conditions. (C) Analysis of Motifs in RpoD binding sites identified from both *in vitro* and *in vivo* ChIP-mini datasets. A traditional ChIP-exo dataset generated under M9 minimal media condition served as a control. (D) Unique RpoD target genes specific to each condition. Genes associated with SPI-1 are highlighted in purple font, while those linked to SPI-2 are indicated in blue font. (E) Volcano plot displaying RpoD differential binding peaks (DBPs) triggered by environmental changes (log_2_ fold change ≤ -1 or ≥ 1, and false discovery rate < 0.05). Genes related to SPI-2 are marked in blue font, and those pertaining to SPI-1 in purple font. Asterisks highlight intragenic (non-regulatory) binding of RpoD, indicating binding occurring downstream of the regulatory region. (F) The transition of RpoD binding sites on effector genes from intragenic to intergenic region in response to environmental changes. (G) The change in RpoD binding intensity for effector genes. “V_peak number” signifies unique RpoD binding sites found *in vivo*, “T_peak number” refers to unique RpoD binding sites sites found *in vitro*, and “B_peak number” represents common RpoD binding sites across both conditions. Negative DBPs are denoted in blue font, and positive DBPs are indicated in red font.

In response to the environmental shift from macrophage extracellular to intracellular conditions, RpoD binding sites for 349 genes disappeared, of which 12 virulence-related genes were identified (Figure 4D and Supplementary Table S4). These genes included the SPI-1 invasion protein *iagB*, the SPI-1 key regulator *hilA*, the HilA-mediated SPI-1 effector *sopA*, and the SPI-3 virulence protein *rhuM* ^60,61^. Additionally, *qseBC* two-component system (TCS) genes that regulate SPI-1 and SPI-2 genes involved in bacterial invasion and survival in host cells were found only under extracellular conditions ^62^. The remaining six genes (*srfABC*, *msgA*, and *rfbKN*) were associated with virulence effectors, virulence proteins, and O-antigen biosynthesis, respectively.

Under macrophage intracellular conditions, 231 genes with *in vivo* unique RpoD binding sites were identified, including 51 genes related to virulence (Figure 4D). Among these virulence-related genes were all SPI-2 and eight SPI-2 effector genes, except for *spiCA*-*ssaDE*, which exhibited RpoD binding sites under both conditions. These virulence-related genes also include four TCS genes encoding the master regulator of the SPI-2 regulon (*ssrAB*) and the PhoPQ-mediated TCS (*pmrAB*) ^63-65^. Additionally, four SPI genes (SPI-1:1, SPI-3:2, and SPI-5:1), eight other virulence-related genes, and PhoPQ-PmrAB linker protein gene (*pmrD*) were exclusively identified within macrophages. To further explore change in RpoD binding intensity in response to environmental shifts, a comparative analysis of differentially binding peaks (DBPs) was performed. A total of 263 positive DBPs and 259 negative DBPs were identified within the 1,539 overlapping binding sites (Figure S10D). A significant increase in RpoD binding intensity was observed upstream of various virulence-related genes, including nine SPI-2 effectors, three SPI-1 genes, two TCSs, one TF, and four virulence proteins (Figure 4E). Conversely, RpoD binding decreased in the intragenic regions of nine virulence-related genes and the intergenic regions of two SPI-3 genes (*fidL* and *marT*). This result also suggests a key finding from a transcriptional regulation standpoint. Under macrophage intracellular conditions, novel and increased RpoD binding was observed in upstream of transcriptional regulatory cascade genes governing *Salmonella* virulence. This cascade includes three positive DBP genes (*phoPQ* and *slyA*) and five *in vivo* unique target genes (*ssrAB*, *pmrAB*, and *pmrD*), which have been reported to activate the expression of SPI-2 effectors and LPS modification genes to enhance survival and virulence (Figure S11) ^65-70^.

Another intriguing finding related to SPI-2 effectors is that RpoD binding peaks were found in the upstream region of 20 out of 29 SPI-2 effector genes under macrophage intracellular conditions (Table 1). Excluding *sseK2*, 19 genes were associated with *in vivo* unique RpoD binding and positive DBPs. According to Jennings et al.^48^, these SPI-2 effector genes were categorized into various functions crucial under host intracellular conditions (Figure S12). Interestingly, differential preferences were observed among effectors within the same category. In the actin cytoskeleton category, RpoD binding was observed upstream of *steC*, while no binding was detected in *spvB*. Additionally, five effectors related in innate immune signaling were associated with RpoD binding, including *sseK1*, *sseK2*, *sseK3*, *gogB*, and *sspH2*. However, no binding was found in *spvCD*, *sspH1*, *gtgA*, *pipA*, and *gogA*. These results suggest the possible existence of a subgroup of SPI-2 effector genes that may play a critical role in adapting to macrophage intracellular conditions.

**Table 1.** RpoD binding sites associated with SPI-2 T3SS effectors.

| Effector | Binding conditions | LFC | Left end | Right end | Peak ID | Interaction Partner(s) |
| --- | --- | --- | --- | --- | --- | --- |
| **SCV membrane dynamics** |  |  |  |  |  |  |
| *sifA* | *in vivo* |  | 1,269,069 | 1,269,118 | Peak_0541 | PLEKHM1, PLEKHM2, GDP-RhoA |
| *sopD2* | *in vivo* |  | 1,012,313 | 1,012,361 | Peak_0410 | Rab7, Rab32* |
| *sseJ* | *in vivo* |  | 1,731,215 | 1,731,266 | Peak_0788 | GTP-RhoA, Cholesterol |
| *gtgE* | both | 2.91 | 1,101,621 | 1,101,670 | Peak_0462 | Rab29*, Rab32*, Rab38* |
| *steA* | *in vivo* |  | 1,680,127 | 1,680,181 | Peak_0759 | PI(4)P |
| *pipB2* | both | 5.47 | 2,948,185 | 2,948,234 | Peak_1322 | Kinesin-1 |
| **Golgi network association** |  |  |  |  |  |  |
| *sseF* | *in vivo* |  | 1,493,233 | 1,493,287 | Peak_0670 | SseG, ACBD3 |
| *sseG* | *in vivo* |  | 1,493,233 | 1,493,287 | Peak_0670 | SseF, ACBD3 |
| **Adaptive immunity** |  |  |  |  |  |  |
| *sseI* | *in vivo* |  | 1,098,254 | 1,098,303 | Peak_0458 | IQGAP1 |
| *steD* | both | 4.89 | 2,285,132 | 2,285,183 | Peak_1037 | mMHCII, MARCH8 |
| **Actin cytoskeleton** |  |  |  |  |  |  |
| *steC* | both | 2.3 | 1,802,216 | 1,802,265 | Peak_0833 | MEK1*, HSP27* |
| *spvB* | ND |  | ND | ND | ND | G-actin* |
| **Innate immune signaling** |  |  |  |  |  |  |
| *sseK1* | both | 1.37 | 4,388,686 | 4,388,737 | Peak_2007 | FADD*, TRADD* |
| *sseK1* | *in vivo* |  | 4,388,877 | 4,388,926 | Peak_2008 | FADD*, TRADD* |
| *sseK2* | *in vivo* |  | 2,282,663 | 2,282,714 | Peak_1031 | ND |
| *sseK2* | both |  | 2,282,772 | 2,282,823 | Peak_1032 | ND |
| *sseK2* | *in vivo* |  | 2,283,038 | 2,283,089 | Peak_1033 | ND |
| *sseK3* | both | 4.2 | 2,094,308 | 2,094,361 | Peak_0990 | TRIM32, TRADD* |
| *gogB* | both | 1.7 | 2,782,050 | 2,782,099 | Peak_1248 | SKP1, FBXO22 |
| *spvC* | ND |  | ND | ND | ND | p-ERK*, p-p38*, p-JNK* |
| *spvD* | ND |  | ND | ND | ND | XPO2 |
| *sspH1* | ND |  | ND | ND | ND | PKN1* |
| *sspH2* | *in vivo* |  | 2,394,972 | 2,395,021 | Peak_1089 | Ubch5-Ubiquitin, SGT1, NOD1* |
| *gtgA* | ND |  | ND | ND | ND | p65*, RelB* |
| *pipA* | ND |  | ND | ND | ND | p65*, RelB* |
| *gogA* | ND |  | ND | ND | ND | p65*, RelB* |
| **Unknown function** |  |  |  |  |  |  |
| *sifB* | both | 5.54 | 1,702,090 | 1,702,144 | Peak_0772 | ND |
| *steB* | *in vivo* |  | 1,729,393 | 1,729,444 | Peak_0785 | ND |
| *sseL* | both | 2.17 | 2,445,959 | 2,446,008 | Peak_1108 | OSBP, Ubiquitin |
| *sseL* | both | 3.16 | 2,446,103 | 2,446,152 | Peak_1109 | OSBP, Ubiquitin |
| *pipB* | *in vivo* |  | 1,135,870 | 1,135,919 | Peak_0482 | ND |
| *srfJ* | ND |  | ND | ND | ND | ND |
| *slrP* | ND |  | ND | ND | ND | ERdj3, TRX1* |

Furthermore, fluctuations in RpoD binding, transitioning from intragenic to intergenic regions, were observed in effector genes. RpoD binding sites of *sifA* and *sseI* shifted from the intragenic region to the TSS after infection (Figure 4F). Additionally, the RpoD binding intensity at the TSS of *sifB* and *sseL* significantly increased, while intragenic RpoD binding either disappeared or decreased in intensity (Figure 4G). Thus, these fluctuations suggest that RpoD may be recruited to the appropriate location for transcription initiation by other transcription factors under macrophage intracellular conditions.

### The fluctuating binding profiles of H-NS and RpoD overlap at the TSS region of virulence genes under macrophage intracellular conditions

As discussed above, we observed fluctuations in H-NS and RpoD binding profiles of *S.* Typhimurium within macrophages. This observation suggests that the reduction in H-NS binding in response to environmental shifts induces an increase in RpoD binding. To explore this premise, we integrated binding profiles of H-NS and RpoD to identify overlapping binding regions associated with virulence (Figure 5A). In regions where H-NS binding was reduced, a significant increase in RpoD binding intensity was observed at the TSS of six SPI-2 effectors (Figure 5B) and a single virulence protein (Figure S13A).

**Figure 5.**
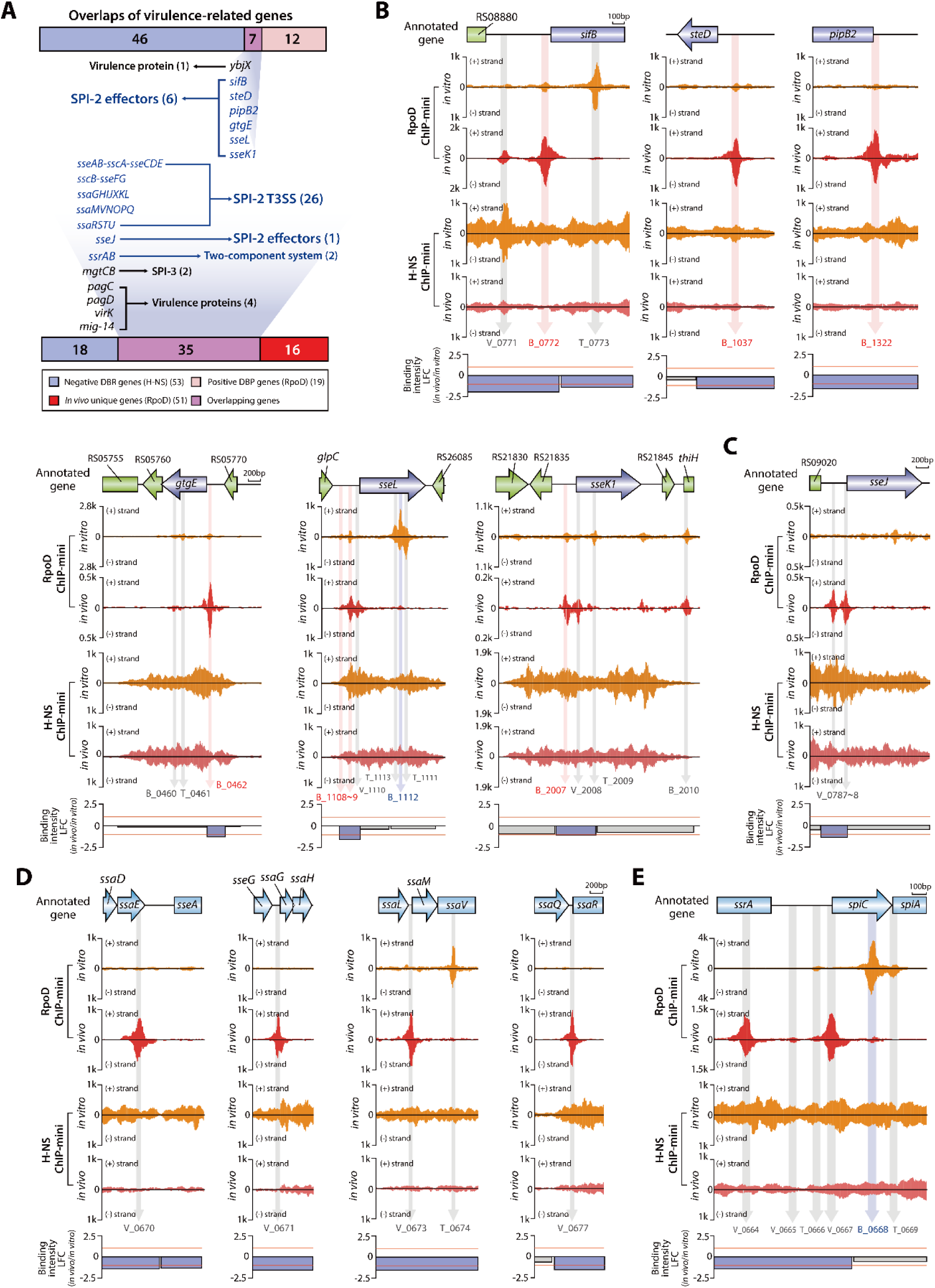
Fluctuations in RpoD binding sites on virulence-associated genes impacted by H-NS negative DBRs. (A) Intersection of virulence-related genes among RpoD target genes and H-NS negative DBR genes in macrophage intracellular conditions. (B) Identification of six effector genes as common elements between RpoD positive DBP genes and H-NS negative DBR genes. (C) The gene *sseJ* recognized as a shared elemnt in *in vivo* unique RpoD genes and H-NS negative DBR genes. (D) Discovery of four *in vivo* unique RpoD binding sites in the SPI-2 region influenced by H-NS negative DBRs. (E) Analysis fo H-NS negative DBR with complex RpoD binding sites in SPI-2. The *spiCA-ssaDE* TU exhibits a single *in vitro* unique RpoD binding site (T_0666) and two *in vivo* unique RpoD binding sites at TSS (V_0665, V_0667).

Furthermore, *in vivo* unique RpoD binding sites upstream of virulence-related genes were identified in H-NS negative DBRs. These binding sites were associated with a single effector gene (*sseJ*) (Figure 5C) and 26 SPI genes (Figure 5D). Notably, four RpoD binding sites in SPI-2 were consistent with previously reported SPI-2 promoter sites ^6^. Moreover, *in vivo* unique binding sites also were detected at the TSS of SPI-3 TU (*mgtCB*) and four virulence protein genes (Figure S13B). The intergenic region of *spiCA-ssaDE* and *ssrAB* showed complex RpoD binding patterns (Figure 5E). Within this complex binding pattern, weak *in vivo* unique binding for *ssrAB* (V_0665) and a shift of binding from the intragenic to intergenic region for *spiCA-ssaDE* were observed, suggesting that RpoD predominantly binds to both TUs under macrophages intracellular conditions. Additionally, strong binding in the *ssrA* intragenic region (V_0664) is associated with *spiCA-ssaDE* ^6^.

To determine a causal relationship between the fluctuation of H-NS and RpoD binding profiles and the alteration in transcript levels, RNA-seq experiments were performed for macrophage extracellular and intracellular *S.* Typhimurium. A total of 1,960 differentially expressed genes (DEGs) (absolute log2 fold change ≥ 1.0 and a false discovery rate < 0.05) were observed by comparing the transcript expression levels of *S.* Typhimurium before and after infection, with 910 genes up-regulated and 1,050 genes down-regulated (Figure S14A and Table S5). Surprisingly, expression of virulence-related RpoD target genes exclusively found inside macrophage showed significant up-regulation (T-test *p*-value < 0.05) (Figure 6A). A decrease of H-NS binding and an increase of RpoD binding were found to led to a notable up-regulation in expression of their target genes (Figure 6B). Additionally, expression of SPI-2 effector genes related in novel or increase of RpoD binding also found to more prominent increases in expression levels than other effector genes (Figure S14B). These findings also highlight our premise that a reduction in H-NS binding within macrophages induces RpoD binding, facilitating the transcription initiation of virulence genes (Figure 6C). Consequently, under macrophage extracellular conditions, we determined that H-NS binds tightly, restricting the access of RpoD for initiation of transcription in virulence-related genes. Conversely, macrophage intracellular conditions lead to a significant reduction in H-NS binding at the TSS region, thereby enhancing the accessibility of RpoD results in novel binding events and increased binding intensity to overcome the silencing effect for transcription initiation (Figure 6D).

**Figure 6.**
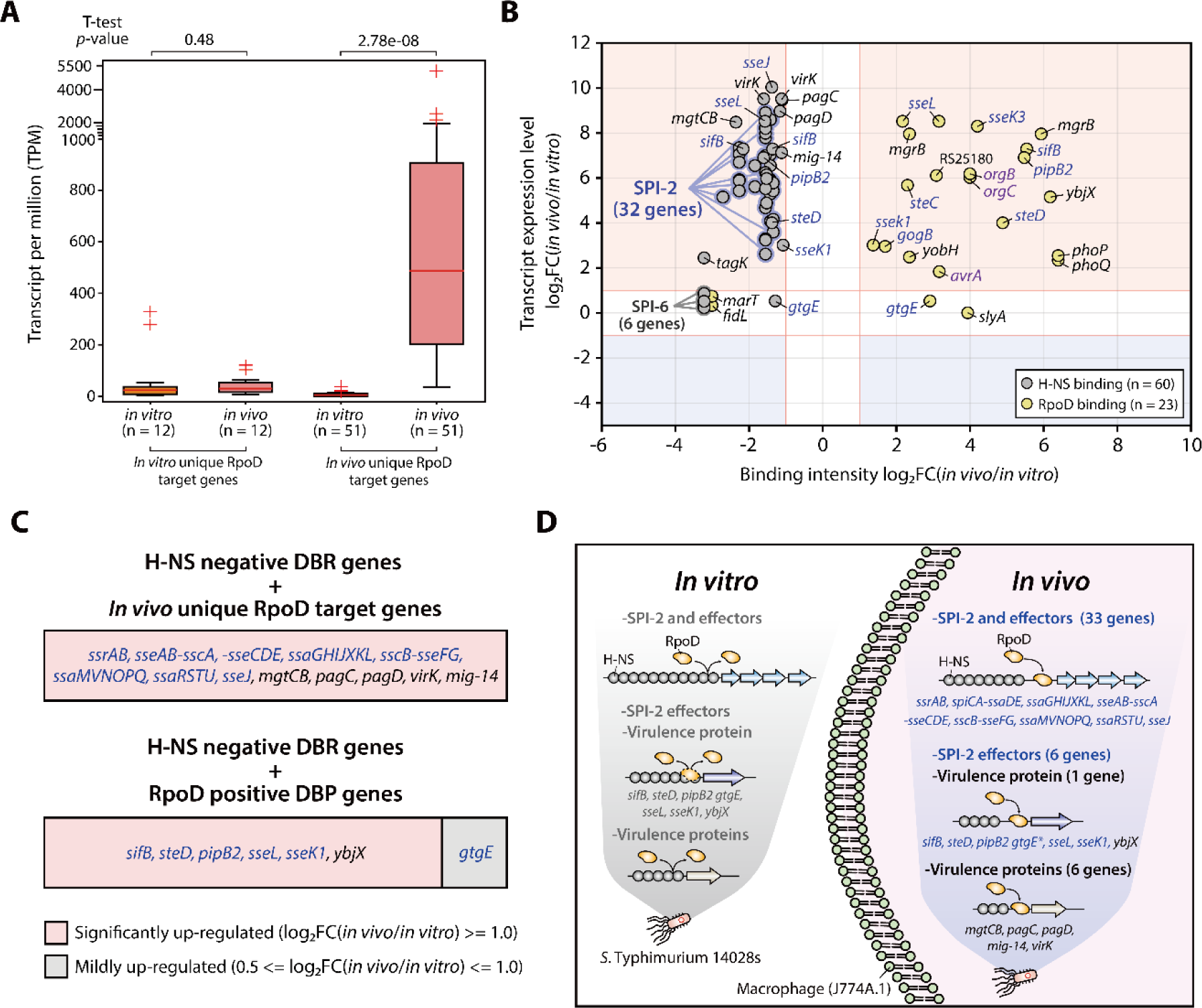
Comprehensive analysis of H-NS and RpoD interaction in regulating transcription initiation of virulence-related genes under macrophage intracellular conditions. (A) Box plots illustrate changes in the expression levels of virulence-related genes corresponding to RpoD binding unique to macrophage extracellular and intracellular conditions. The expression of RpoD target genes uniquely identified within macrophage was found to be significantly up-regulated (t-test *p*-value < 0.05). (B) Scatter plot depicts the correlation between change in binding intensities of H-NS and RpoD with corresponding virulence-related gene expression levels in response to environmental shift. Genes related to SPI-2 are marked in blue, and those pertaining to SPI-1 in purple. (C) An increase in RpoD binding at H-NS negative DBRs induces transcription initiation of virulence-related genes, consequently leading to a significant up-regulation in their transcriptional expression. (D) Overview highlights how the binding dynamics of H-NS and RpoD on virulence-related genes in *S.* Typhimurium 14028s modulate transcription initiation in response to the environmental transition from macrophage extracellular to intracellular conditions.

## Discussion

In this work, we optimized the traditional ChIP-exo method to overcome previous challenges in identifying binding profiles of DNA-binding proteins with a near single-base pair resolution for host intracellular bacteria. ChIP-mini clearly surpasses other ChIP-based methods in identifying binding profiles at genome-wide levels, even when using a minimal initial quantity of bacterial cells (4.8×10^6^). Furthermore, ChIP-mini shows a nearly complete overlap with traditional ChIP-exo binding profiles, when tested with *E. coli* RpoD (>83%).

Despite the importance of understanding TRNs during infection, comparative analysis of genome-wide binding profiles between host extracellular and intracellular conditions has been challenging due to the use of a low quantity of bacterial cells in infection experiments. This limitation necessitated the use of ChIP-mini to directly measure the impact of the environmental shifts on binding of DNA-binding proteins with improved resolution. For *S.* Typhimurium, a ubiquitous and clinically significant intestinal pathogen, we utilized ChIP-mini applications and the DiffExo pipeline to reveal the crucial interaction between H-NS and RpoD concerning virulence genes during macrophage infection. The reduction in H-NS binding in intergenic regions exposes the promoter, leading to 1) binding of RpoD to the promoter of SPI-2, SPI-2 effector, SPI-3, and virulence protein genes, and 2) an increase in RpoD binding to SPI-2 effectors and virulence protein genes (Figure 6D). Remarkably, these findings consistent with significantly up-regulated expression levels of those genes, highlighting a crucial regulatory mechanism whereby diminished H-NS binding enables enhanced RpoD binding to induce transcription initiation of key virulence genes after infection. By doing so, *S.* Typhimurium ensures the effective activation of transcription initiation of virulence genes for proliferation and replication when it is inside host cells. In addition to virulence-related genes, ChIP-mini data identified RpoD target genes exhibiting increased binding inside macrophages, which were implicated in various metabolic processes (Figure 4D). Thus, our integrated ChIP-mini data and transcription profiles not only provide genome-wide binding information of H-NS and RpoD, but also offer a valuable resource for future studies to elucidate complex metabolic perturbations in *S.* Typhimurium during host infection.

Additionally, we identify a transcriptional regulator, PhoP, with increased RpoD binding and transcript expression within macrophages. This regulator is known to alleviate H-NS silencing and recruit RNA polymerase to horizontally acquired genes ^71^. Notably, activated PhoP binding sites were mainly found upstream of the RpoD binding sites in macrophage intracellular *S.* Typhimurium, acting as transcriptional activators (Figure S15A). Moreover, the majority of PhoP regulon genes were also identified among *in vivo* RpoD target genes (Figure S15B). This observation suggests a complex regulatory network involving RpoD and virulence transcriptional regulators such as PhoP, SlyA, HilD, SsrB, and OmpR, which proactively interact with H-NS binding regions within macrophages ^37,71,72^. Thus, future efforts to reveal genome-wide interactions of these transcriptional regulators with H-NS and RpoD using ChIP-mini applications would greatly expand our understanding of the comprehensive regulatory networks responding to macrophage intracellular conditions.

Using ChIP-mini, we are able to identify the binding dynamics of H-NS and RpoD with a near-base pair resolution during the transition from macrophage extracellular and intracellular conditions, highlighting the crucial interaction between H-NS and RpoD at transcriptional initiation standpoint. Incidentally, we confirmed the feasibility of adapting ChIP-mini applications to other bacteria and host cells, successfully identifying genome-wide binding sites of HrcA in *Chlamydia trachomatis* infected mouse fibroblast L929 cells. Hence, the ChIP-mini workflow will be a powerful tool to expedite the study of TRN reconstruction in wide range of host intracellular pathogens, requiring only minor modifications to the experimental process described herein.

## STAR★Methods

### Key Resources Table

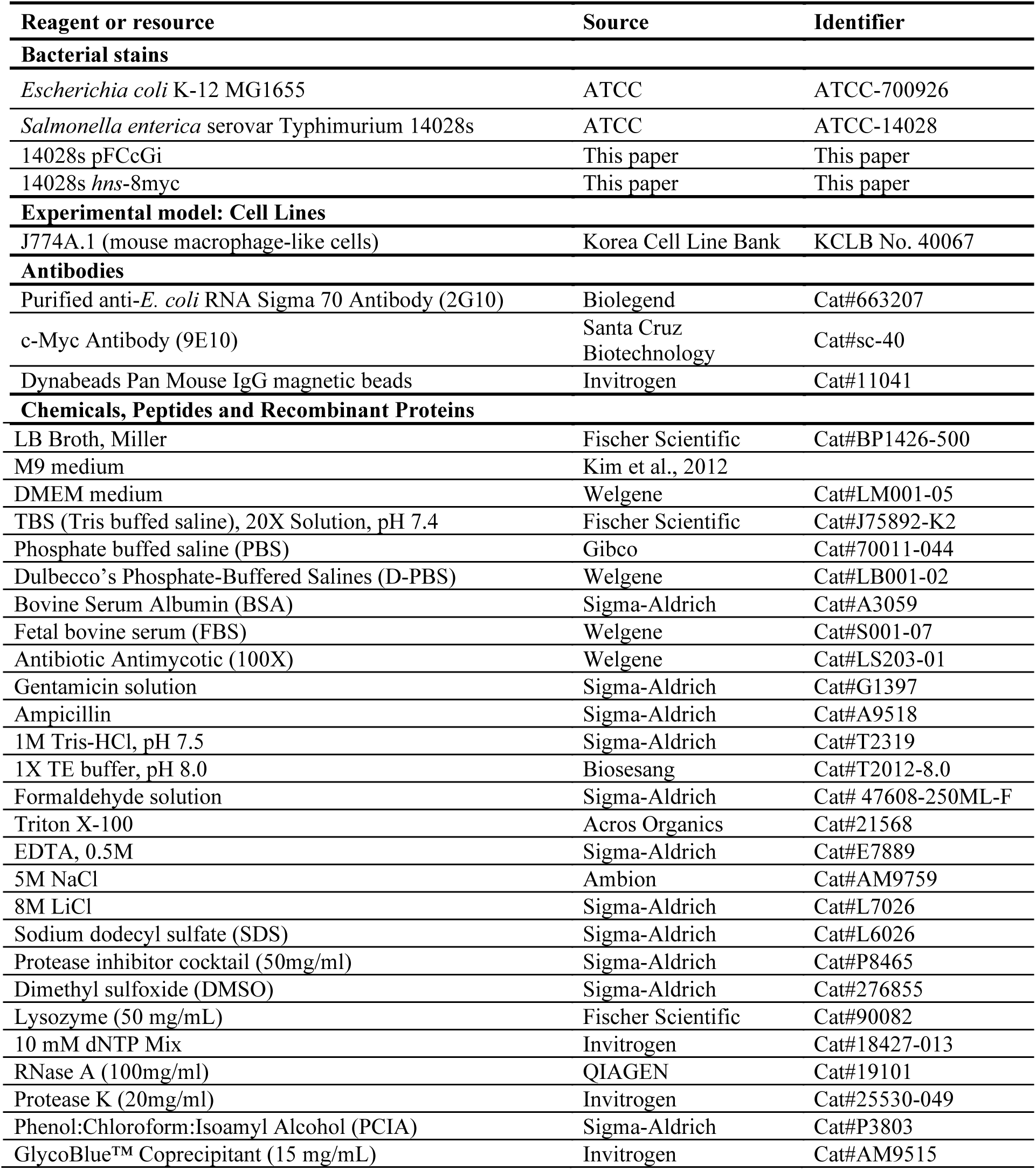
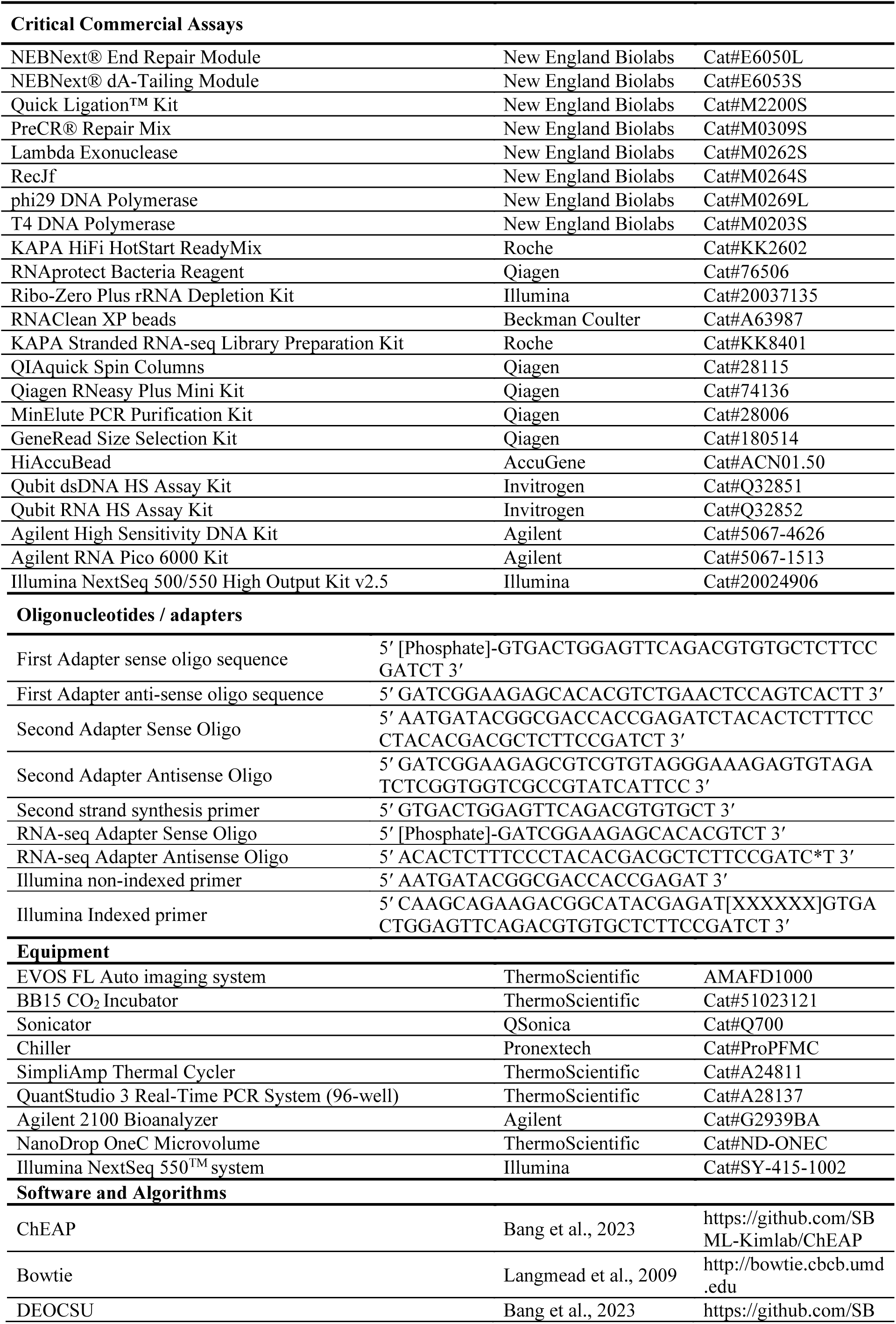
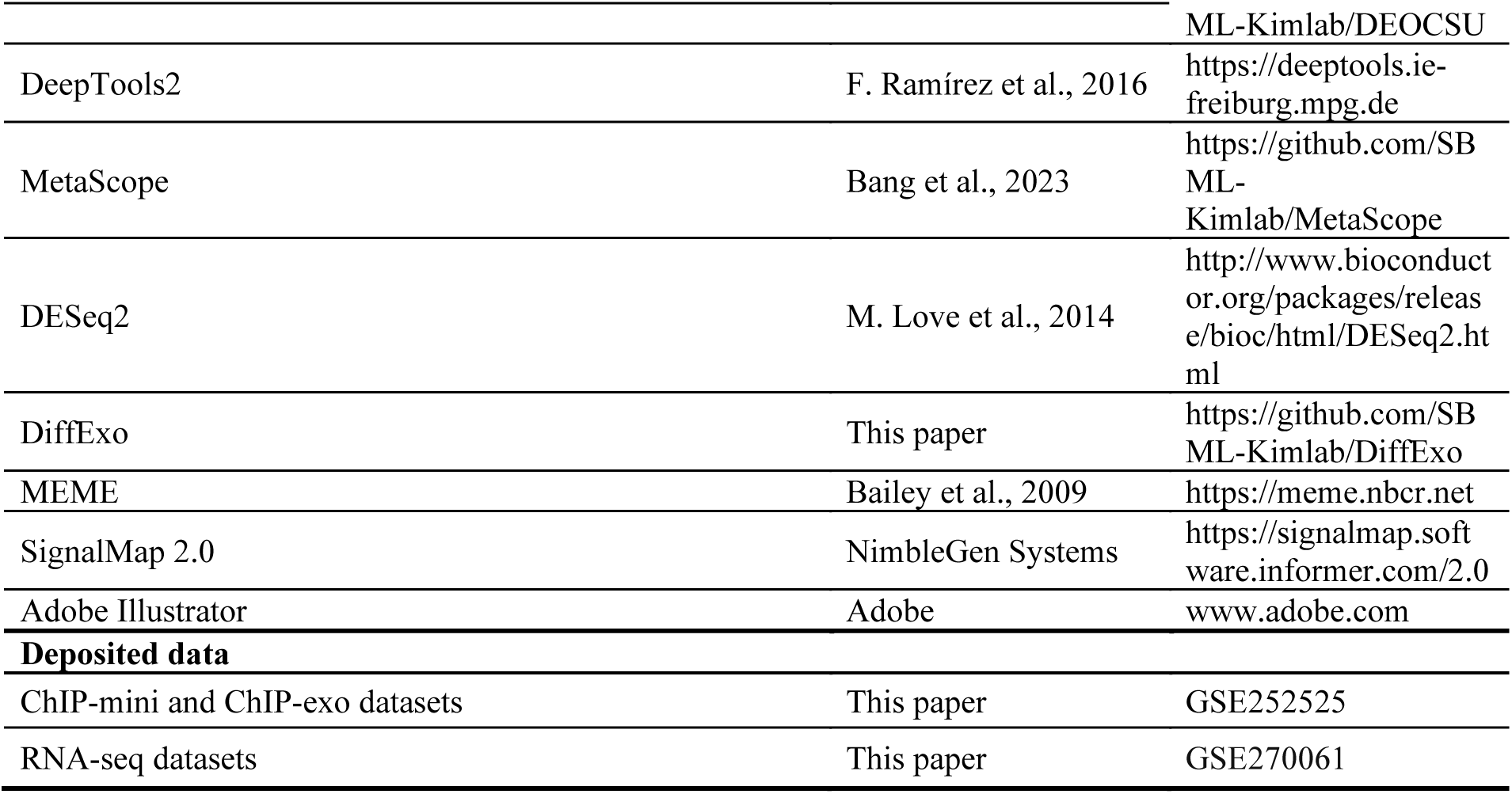

### Experimental Model and Subject Details

#### Bacterial strains and growth conditions

All bacterial strains described in this study listed in the Key Resources Table. For ChIP-mini applications, *S.* Typhimurium harboring *hns*-8myc was generated through a λ red-mediated site-specific recombination system as described previously ^73^. For ChIP-exo and ChIP-mini experiments, glycerol stock of *E. coli* K-12 MG1655 was inoculated into M9 minimal media with 0.2% (w/v) glucose. M9 minimal media was also supplemented with 1 ml trace element solution (100x) containing 1 g EDTA, 29 mg ZnSO_4_·7H_2_O, 198 mg MnCl_2_·4H_2_O, 254 mg CoCl_2_·6H_2_O, 13.4 mg CuCl_2_ and 147 mg CaCl_2_. The inoculated culture was incubated at 37 °C overnight with constant agitation. Culture was then diluted into 100 ml of fresh minimal media (1/200 dilution) and cultured at 37 °C with agitation in a water bath to the mid-exponential phase (OD_600_ ≈ 0.6∼0.7). For *in vitro* and *in vivo* ChIP-mini experiments, glycerol stocks of *S.* Typhimurium wild-type and *hns*-8myc tagged strains were inoculated into LB broth. Cultures were incubated at 37 °C overnight with agitation, and then was used to inoculate the 2ml of fresh LB broth. The fresh culture was incubated at 37 °C overnight with agitation in a shaking incubator. For fluorescent image analysis measuring *Salmonella* infection efficiency, glycerol stock of *S.* Typhimurium harboring pFCcGi vector was inoculated to LB broth with 100 μM ampicillin and incubated at 37 ℃ overnight with constant agitation. The culture was then diluted into 2ml of fresh LB broth with 100 μM ampicillin and cultured at 37 °C overnight with agitation in a shaking incubator.

#### Cell culture

The macrophage-like cell line (J774A.1) cells employed in this study was cultured in Dulbecco’s modified Eagle’s medium (DMEM) supplemented with 10% (v/v) fetal bovine serum (FBS) and antibiotic antimycotic at 37 °C with 5% CO^2^ in a humidified incubator. J774A.1 cells were seeded in 6-well plates at a plating density of 5×10^5^ per well for fluorescent image analysis. For ChIP-mini applications, J774A.1 cells were seeded in 75T flasks at a plating density of 6 x 10^6^ per flask.

### Method Details

#### Fluorescent image analysis of *Salmonella* infection

The macrophage cells were seeded in 6-well plates at a plating density of 5×10^5^ per well under the culture conditions described above. Overnight-grown *S.* Typhimurium harboring pFCcGi vector in LB broth with 100 μM ampicillin were added to the macrophages at a multiplicity of infection (MOI) of 10 ^44,45^. The plates were centrifuged at 500 xg for 5 minutes at room temperature and incubated for an additional 30 minutes. To remove extracellular bacteria, two rounds of phosphate-buffered saline (PBS) washing were conducted, followed by DMEM supplemented with 10% (v/v) FBS and 150 ug ml^-1^ gentamicin. After 1hour, the DMEM was replaced to fresh DMEM containing 15 μg ml^-1^ gentamicin, and the plates were then incubated for 24 hours at 37 ℃ with 5% CO^2^ in a humidified incubator. Fluorescent images were obtained using EVOS FL Auto imaging system.

#### ChIP-exo experiment

ChIP-exo for bacteria was performed following the procedures described in previous studies ^22-24^. To identify RpoD binding profiles, 50 ml of initial *E. coli* culture (number of cells = 2.4×10^10^) was used to crosslink DNA and RpoD using formaldehyde at mid-exponential phase. Crosslinked cells were resuspended in 270.5 μl of lysis buffer (10 mM Tris-HCl, pH 7.5, 100 mM NaCl, and 1 mM EDTA) including protease inhibitor cocktail and lysozyme and diluted in 275 μl of IP buffer (100 mM Tris-HCl, pH 7.5, 200 mM NaCl, 1 mM EDTA, and 2% Triton X-100 (w/v)) to lyse cells and fragment DNA using sonication (25 minutes with 50s on and 10s off intervals and amplitude 50%). Subsequently, we isolated the DNA bound to RpoD from the crosslinked *E. coli* lysate by chromatin immunoprecipitation (ChIP) with the specific antibody recognizing RpoD (Anti-RpoD mouse IgG) and Dynabeads Pan Mouse IgG magnetic beads. This process was followed by stringent washings ^13^.

ChIP materials (chromatin-Dynabeads) were used to perform on-bead enzymatic reactions according to the ChIP-exo method ^21^. In Stage 1, the sheared DNA on Dynabeads was repaired using 100 μl of the NEBNext End Repair Module. 50 μl of the dA-Tailing Module added a single dA overhang, followed by ligation of the first adaptor (5′-phosphorylated) using 50 μl of the NEBNext Quick Ligation Module. Nick repair was performed using 50 μl of PreCR Repair Mix. DNA was treated with 50 μl of Lambda exonuclease Mix and RecJf exonuclease Mix to digest the 5’ ends of DNA. Exonuclease-treated chromatin was eluted from the beads using 200 μl of elution buffer (50 mM Tris-HCl, pH 8.0, 1 mM EDTA, and 1% SDS (w/v)) by overnight incubation at 65 °C, and 200 μl of TE buffer was added to make up the proper volume for Phenol:Chloroform:Isoamyl Alcohol (PCIA) purification and ethanol precipitation. To deplete RNA and reverse the protein–DNA crosslink, RNaseA and Proteinase K were used, respectively.

In Stage 2, RNAs and proteins-removed samples purified by PCIA/ethanol precipitation were used to perform primer extension and 2^nd^ adapter ligation with the following modifications. The DNA samples were incubated for primer extension of 2^nd^ strand synthesis as described previously ^22^, followed by dA tailing using 8 μl of the dA-Tailing Module and second adaptor ligation using 27 μl of the NEBNext Quick Ligation Module. To remove 3’ overhangs, 30 μl of T4 DNA polymerase Mix was used. The DNA samples from each step in Stage 2 were purified by QIAquick or MinElute Spin Columns. Finally, the DNA samples purified by GeneRead Size Selection Kit were enriched by polymerase chain reaction (PCR) using KAPA HiFi HotStart ReadyMix. The amplified DNA samples were purified again by GeneRead Size Selection Kit and quantified using Qubit dsDNA HS Assay Kit. Traditional ChIP-exo experiments were performed in biological duplicate.

#### ChIP-mini experiment

A detailed, ChIP-mini method depending on number of initial bacterial cells is presented in Supplementary Methods. In the minimalized ChIP-mini method, crosslinked cells were resuspended in 108.2 μl of lysis buffer mix, protease inhibitor cocktail, and lysozyme. 110 μl of the IP buffer was added to lyse cells. DNA for RpoD samples were fragmented using sonication (40 minutes with 50s on and 10s off intervals and amplitude 50%). ChIP was proceeded using 1.5 μl of RpoD antibody or 3.75 μl of c-Myc antibody, and 15 μl of Dynabeads Pan Mouse IgG magnetic beads followed by stringent washings. In Stage 1, the sheared DNA on Dynabeads was repaired using 10 μl of the NEBNext End Repair Module, followed by 5 μl of the dA tailing Module, 1^st^ adaptor ligation using NEBNext Quick Ligation Module, Nick repair Mix, Lambda exonuclease Mix, and RecJf exonuclease Mix. Exonuclease-treated chromatin was eluted from the beads using 20 μl elution buffer by overnight incubation at 65 °C. RNaseA and Proteinase K were used to remove RNAs and to reverse-crosslink chromatin, respectively. These RNAs and proteins-removed samples (>100 bp) were purified by 2.5x HiAccuBead.

In Stage 2, we developed a one-tube procedure for DNA purification to minimize the loss of DNA samples. This procedure allows the use of an enzyme mixture from each step as a bead-elution buffer, introducing the beads only once and adjusting the polyethylene glycol (PEG) concentration to purify DNA of the appropriate size generated at each step. The DNA samples were incubated for primer extension of 2^nd^ strand synthesis as described previously ^22^. DNA with the second strand synthesized was purified by 2.5x HiAccuBead and eluted in 10 μl elution buffer consisting of 1.6 μl dA-Tailing Module and 8.4 μl filtered DW. After dA-tailing, the enzyme was deactivated at 60 °C for 30 minutes, followed by the addition of 15 μl of the NEBNext Quick Ligation Module for the 2^nd^ adapter ligation. This adapter-ligated DNA with beads was purified by adding 25 μl Polyethylene glycol (PEG). Subsequently, the 3’ overhang was removed using 10 μl of the T4 DNA polymerase Mix as the elution buffer. This DNA sample with beads underwent purification by adding 10 μl of PEG, followed by elution with 20 μl of filtered DW. Finally, the DNA sample was enriched by PCR using KAPA HiFi HotStart ReadyMix. The amplified DNA samples were purified again by 1.0x HiAccuBead and quantified using Qubit dsDNA HS Assay Kit. ChIP-mini experiments were also performed in biological duplicate.

#### *In vitro* and *in vivo* ChIP-mini experiments

Both ChIP-mini applications were conducted using a macrophage with *S.* Typhimurium wild-type or *hns*-8myc tagged strains. The macrophage cells were seeded in 75T flasks at a plating density of 6×10^6^ per flask under the culture conditions described above. Overnight-grown bacteria were added to the macrophages at a multiplicity of infection (MOI) of 10, and the 75T flasks were centrifuged at 500 xg for 5 minutes at room temperature and incubated for an additional 30 minutes ^44,45^. Following infection, extracellular bacteria in DMEM were isolated and crosslinked using formaldehyde for ChIP-mini (*in vitro* Chip-mini). The remaining extracellular bacteria were washed three times with PBS and killed by 1 hour incubation with DMEM supplemented with 10% FBS and 150 μg ml^−1^ gentamycin. For post-infection, the DMEM was replaced to fresh DMEM containing 10% FBS with 15 μg mL^−1^ gentamicin, and the flasks were incubated at 37 °C for 6 hours. After 6 hours incubation, *S.* Typhimurium-infected macrophages were co-crosslinked using formaldehyde. Co-crosslinked cells were lysed with 1% Triton X-100 and centrifuged at 20,000xg for 5 minutes at 4 °C. The cell pellet containing intracellular bacteria was utilized for the ChIP-mini method (*in vivo* Chip-mini). A detailed, ChIP-mini application experiments is presented in Supplementary Methods.

#### *In vitro* and *in vivo* RNA-seq experiments

Extracellular and intracellular *S.* Typhimurium wild-type cells from macrophages were cultured under identical growth conditions as described for the ChIP-mini applications. For total RNA isolation of extracellular bacteria, 10 ml DMEM containing extracellular *S.* Typhimurium after infection were mixed with 30 ml Qiagen RNAprotect Bacteria Reagent. Samples were immediately mixed by vertexing for 5 seconds, incubated at room temperature for 5 minutes, and then centrifuged at 5,000xg for 10 minutes. The supernatant was decanted, any residual supernatant was removed by inverting the tube once onto a paper towel. For total RNA isolation of intracellular bacteria, *S.* Typhimurium-infected macrophages were mixed with 30 ml of Qiagen RNAprotect Bacteria Reagent and incubated at room temperature for 5 minutes. After decanting the supernatant, the infected macrophages were transferred to 1.7 mL microtube and subsequently centrifuged at 20,000 xg for 5 minutes. Total RNA samples were then isolated using the Qiagen RNeasy Plus Mini kit according to the manufacturer’s instructions. The samples were quantified using a NanoDrop 1000 spectrophotometer, and the quality of the isolated RNA was assessed with an RNA 6000 Pico Kit on an Agilent 2100 Bioanalyzer. Ribosomal RNA depletion was subsequently carried out with the Illumina Ribo-Zero Plus rRNA Depletion Kit. Following rRNA depletion, a paired-end, strand-specific RNA-seq library was constructed using the KAPA Stranded RNA-seq Library Preparation Kit according to the manufacturer’s instructions. All RNA-seq experiments were performed in biological duplicate.

#### Sequencing and Data analysis

Quality of the DNA samples were checked by running an Agilent High Sensitivity DNA Kit using Agilent 2100 Bioanalyzer before sequencing. Paired-end sequencing (41 bp reads) was performed on the Illumina NextSeq550 using Illumina TruSeq Read_1 and Read_2 primers. For ChIP-mini and traditional ChIP-exo data analysis, sequence reads were mapped to the reference genomes of *E. coli* K-12 MG1655 (NC_000913.3) or *S.* Typhimurium 14028s (NC_016855.1 and NC_016856.1) by bowtie with default options to generate SAM output files using ChEAP (ChIP-exo analysis pipeline), these output files were converted to BAM and GFF files for further analysis ^74^. DEOCSU (DEep-learning Optimized ChIP-exo peak calling Suite, https://github.com/SBML-Kimlab/DEOCSU) ^27^, a novel machine learning-based ChIP-exo peak-calling suite pipeline, was used to define binding peak candidates from the biological duplicates. To reduce false-positive peaks, those with a signal-to-noise ratio (S/N) less than 1.0 were removed. DeepTools2 was used to compute the correlation of sequencing libraries with the option of BAM coverage calculation to consecutive bins of equal size (10 bp) ^25^. Genome-scale data were visualized using GFF files via MetaScope (https://github.com/SBML-Kimlab/MetaScope) ^27^. For RNA-seq analysis, sequence reads were mapped onto the reference genome (NC_016855.1 and NC_016856.1) using bowtie ^75^ with the maximum insert size of 1,000 bp, and 2 maximum mismatches after trimming 3 bp at 3’ ends to generate SAM files. These output files were also converted to BAM and GFF files for further analysis. To compare expression changes according to environmental shift from macrophage extracellular to intracellular conditions, featureCount and DESeq2 was used to calculate transcripts per million (TPM) value and differential expression ^31^. From DESeq2 output, genes with differential expression with absolute value of log2 fold change ≥1.0 and false discovery rate <0.05 were considered differentially expressed genes (DEGs).

#### DiffExo: ChIP-exo peak normalization pipeline

The DiffExo pipeline, comprising four Python and one R scripts, was developed to perform the analyses outlined in this study (Figure S3) (https://github.com/SBML-Kimlab/DiffExo) ^30,31^. The RPPM (Reads Per Peak per Million) value was conceived to normalize ChIP-exo and ChIP-mini sequencing datasets. The calculation of RPPM for peak *i* uses the following formula:

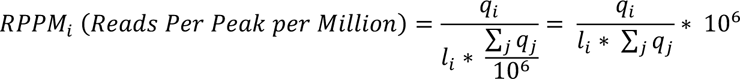

where *q*_*i*_ are raw sequencing read counts in peak *i*, *l*_*i*_ is length of ChIP-exo or ChIP-mini binding peak, and ∑_*j*_ *q*_*j*_ corresponds to the total number of mapping reads on the reference genome. Statistically, significant differentially binding sites (DBRs and DBPs) were calculated using DESeq2, which adopted a negative binomial distribution-based model ^31^. From the DESeq2 output, peaks with absolute value of log_2_ fold change ≥ 1.0 and false discovery rate < 0.05 were considered as differentially binding sites.

#### Motif analysis from the ChIP-exo and ChIP-mini RpoD binding peaks

The RpoD binding motif analyses were completed using MEME tools from the MEME software suite ^76^. Sequences were extended by 20 bp from target genes, because only -10 box, gntAtaaT, was found without that extension. This observation conflicts with the knowledge of RpoD, because RpoD is known to specifically recognize -10 and -35 box sequences, which are expected to be covered and protected from exonuclease activity. RpoD binding peaks align well with RpoB binding peaks (Figure S16), and TSSs were dominantly located at the center of binding regions; thus ChIP-exo might be capturing RpoB bindings that were associated with RpoD ^15^.

## Supporting information

Table S1_Total_binding sites of H-NS and RpoD

Table S2_Differentially binding regions (DBRs) and differentially binding peaks (DBPs)

Table S3_Target genes of differentially binding regions and peaks

Table S4_RpoD unique target genes under macrophage in vitro and in vivo conditions

Table S5_Differentially expressed genes (DEGs) due to enviromental conditions

## Data Availability

The whole dataset of ChIP-mini and RNA-seq has been deposited to GEO with the accession number of GSE252525 and GSE270061, respectively. DiffExo pipeline is freely available at the public GitHub repository (https://github.com/SBML-Kimlab/DiffExo).

## Acknowledgement

This research was supported by National Research Foundation of Korea (NRF) funded by the Ministry of Science and ICT (MSIT) [2021M3A9I402484011, 2022M3A9I5018934]; the UNIST Center for Waste Plastics Carbon Cycling (UWCC) funded by The Circle Foundation, Republic of Korea.

## Author contributions

JYP, MJ, and DK conceived the study. JYP and MJ performed all traditional ChIP-exo, ChIP-mini, RNA-seq experiments. JYP and JW performed the computational analysis and constructed the DiffExo pipeline. EC and SK performed the macrophage infection experiments. DK and EJL supervised the study. IN operated the Illumina NextSeq500 for sequencing. JYP, MJ, SML, EJL, and DK wrote the manuscript. All authors helped edit the final manuscript. JYP and MJ contributed equally to this work.

