## Supplementary Information for "Elucidating DNA-binding protein dynamics in *Salmonella* Typhimurium within macrophages using a breakthrough low-input ChIP-exo approach"

**Supplementary Text S1**

To generate a ChIP-exo library from a low number of bacterial cells, ChIP-mini was optimized based on the traditional ChIP-exo method [1-4]. Traditional ChIP-exo and ChIP-mini share the same procedures consisting of four parts as follows: 1) DNA fragmentation, 2) exonuclease digestion of antibody-TF complex (Stage 1), 3) reverse cross-linking, and 4) construction of sequencing library (Stage 2).

In the sonication procedure, we optimized both the volume of the fragmentation buffer and the sonication time for different numbers of crosslinked *E. coli* cells (Figure 1C). The $3.0x10^9$ and $7.68x10^8$ samples were fragmented by 25 minutes of sonication, utilizing 50% and 40% of the total volume of fragmentation buffer required by traditional ChIP-exo, respectively. For the remaining four samples, 40 minutes of sonication with 40% of the total volume of the fragmentation buffer, as compared to the traditional method, was used. It was observed that the efficiency of sonication decreased when less than 30% of the total buffer volume was used. Given that traditional ChIP-exo for bacteria typically utilizes DNA fragments with starting lengths ranging from 200 to 600 bp, this modification enabled targeted fragmentation even in small numbers of bacterial cells (Figure S2B). Following the reduction of the fragmentation buffer, reagents for the subsequent steps in ChIP-mini were optimized to minimize volume. The volumes of 1st antibody and Dynabeads were minimized to 25% to avoid non-specific binding of antibodies, and reagents required for Stage 1 including lambda exonuclease digestion and 1st adaptor ligation were reduced to 10% compared to ChIP-exo (Figure S1C). For the $3.0x10^9$ sample, the volumes of 1st antibody, Dynabeads, and Stage 1 reagents were reduced to 50%, 50%, and 25%, respectively compared to ChIP-exo.

In the case of ChIP-seq, the reverse cross-linking step involving DNA purification is known to be one of the key steps resulting in immunoprecipitated-DNA (IP-DNA) loss [5]. ChIP-mini is more susceptible to IP-DNA loss caused by reverse cross-linking because it uses a minimal number of bacterial cells. Furthermore, this loss of IP-DNA can further hinder not only Stage 2 procedures but also generate amplification bias due to the small amount of initial adapter-ligated DNA. Thus, it was considered a crucial point to maintain the efficiency of IP-DNA purification when using reduced volume of the reverse cross-linking solution. According to traditional ChIP-exo, Phenol:Chloroform:Isoamyl Alcohol (PCIA) and ethanol precipitation methods are used to purify the IP-DNA after the reverse cross-linking step. These two experimental procedures are difficult to perform on microtube scale volumes and consume a relatively long time, over a couple of hours. To mitigate these disadvantages, a bead-based purification method that can be used on small volumes and takes a short time (< 20 min) was verified as an alternative, which was also adopted in ChIP-exo 5.0 for eukaryotes [6].

To verify the validity of this method, we used traditional ChIP-exo samples containing anti-RpoD antibodies to measure the concentration of IP-DNA extracted via PCIA or bead-based methods after reverse cross-linking and 2nd strand synthesis steps. After the reverse cross-linking step, one sample was divided into two duplicates to be purified using the PCIA/ethanol precipitation method or 2.5x DNA purification beads method with a high binding affinity for 1st adapter-ligated IP-DNA longer than 100bp. As a result, 0.304 ng and 0.342 ng of IP-DNA were conserved using PCIA/ethanol precipitation and bead-based purification, respectively. This amount of IP-DNA from both methods showed that the bead-based method has slightly higher efficiency for DNA purification than the PCIA/ethanol precipitation method from traditional ChIP-exo. Thus, these results suggest that bead-based purification can maintain efficiency compared to PCIA-based purification, even with a low volume of the reverse cross-linking solution.

Finally, to optimize the experimental material of Stage 2 steps for constructing the sequencing library, ChIP-mini also replaces spin column-based DNA purification methods with bead-based DNA purification methods. Since the minimum elution volume of the spin column was 10 μl, the reduction of experimental materials of Stage 2 is limited. Therefore, we developed a one-tube procedure for DNA purification to minimize the loss of DNA samples by using a bead-based purification. This procedure allows the use of an enzyme mixture from each step as a bead-elution buffer, enabling an average reduction of 52% in the volume of reagents required compared to traditional ChIP-exo. Additionally, we introduced the beads only once and adjusted the polyethylene glycol (PEG) concentration to purify IP-DNA of the appropriate size generated at each step. Furthermore, the bead cleanup procedure between the dA-tailing and the 2nd adapter ligation was omitted by using enzyme heat inactivation [7]. This simplified method is anticipated to minimize IP-DNA loss relative to column-based methods. Consequently, the number of PCR cycles needed for library amplification showed no significant difference between the $9.6x10^7$ ChIP-mini sample and the traditional ChIP-exo sample using $2.4x10^{10}$ bacterial cells (Table 1).

**Supplementary Figures**

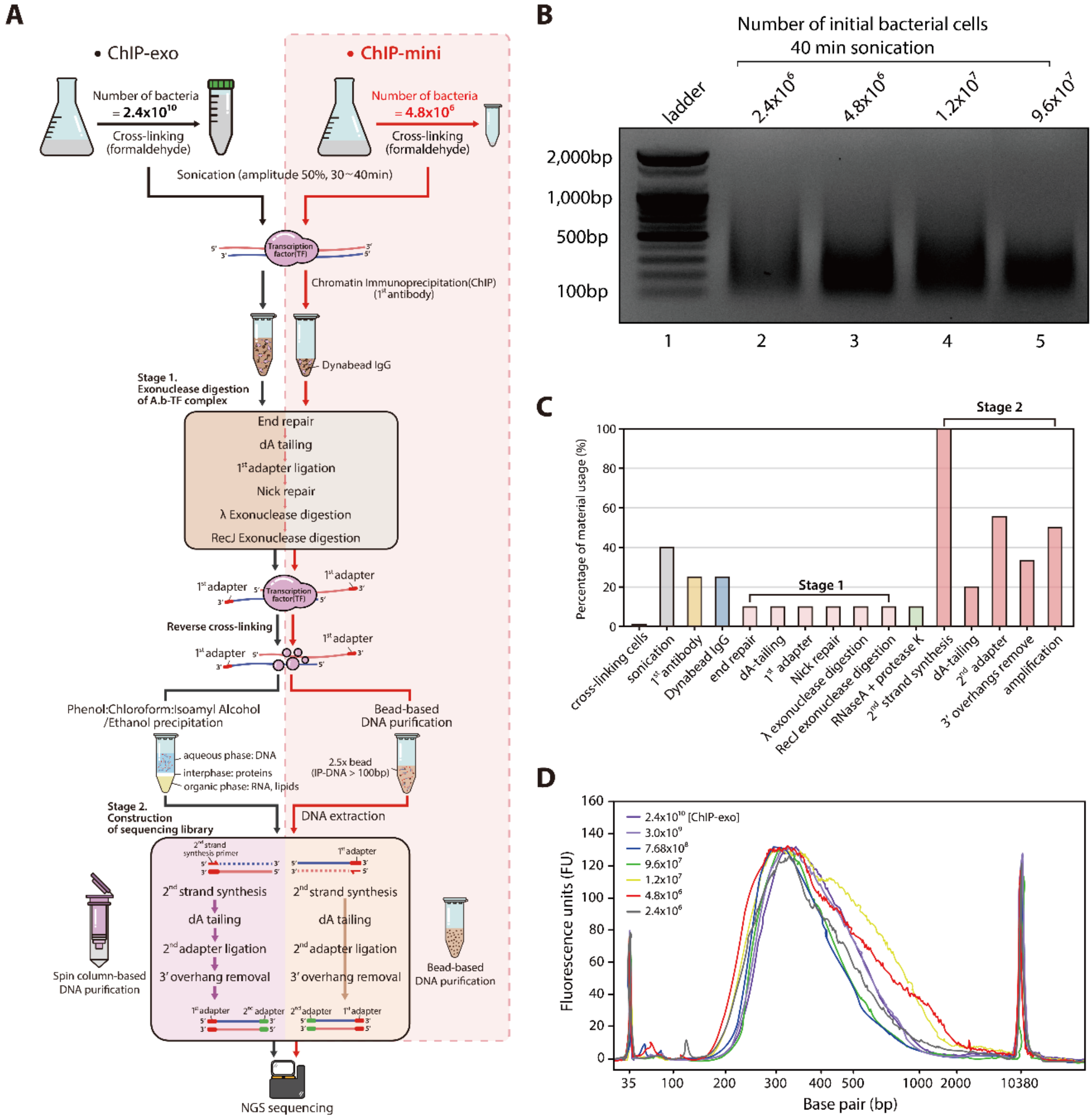

**Figure S1. Comparison of the traditional ChIP-exo and optimized ChIP-mini processes.** (A) Comparison of steps to generate ChIP-exo and ChIP-mini libraries. Different numbers of initial bacterial cells are lysed and sonicated to perform ChIP. The optimized volume of the ChIP process enables minimization of reagents for 1st adapter ligation and exonuclease digestion of IP-DNA including antibody-TF complex (Stage 1). After reverse cross-linking, ChIP-mini uses 2.5x AMPure beads to extract IP-DNA over 100 bp. Additionally, ChIP-mini utilizes a bead-based DNA purification rather than a spin column-based one, allowing for reduced reagent volumes during sequencing library construction (Stage 2). (B) The distribution of DNA fragment sizes based on a low number of bacterial cells in 1% agarose gel after 40 minutes of sonication (< $9.6 \times 10^7$). Sonicated DNA was fragmented to the proper size to proceed with ChIP-mini (200~600 bp). (C) Detailed experimental material usage for the minimum number of cross-linked cells in ChIP-mini. (D) Distribution of ChIP-exo and ChIP-mini libraries for *E. coli* RpoD after PCR amplification. The size distribution of ChIP-mini libraries was similar to that of the libraries generated with the original ChIP-exo method.

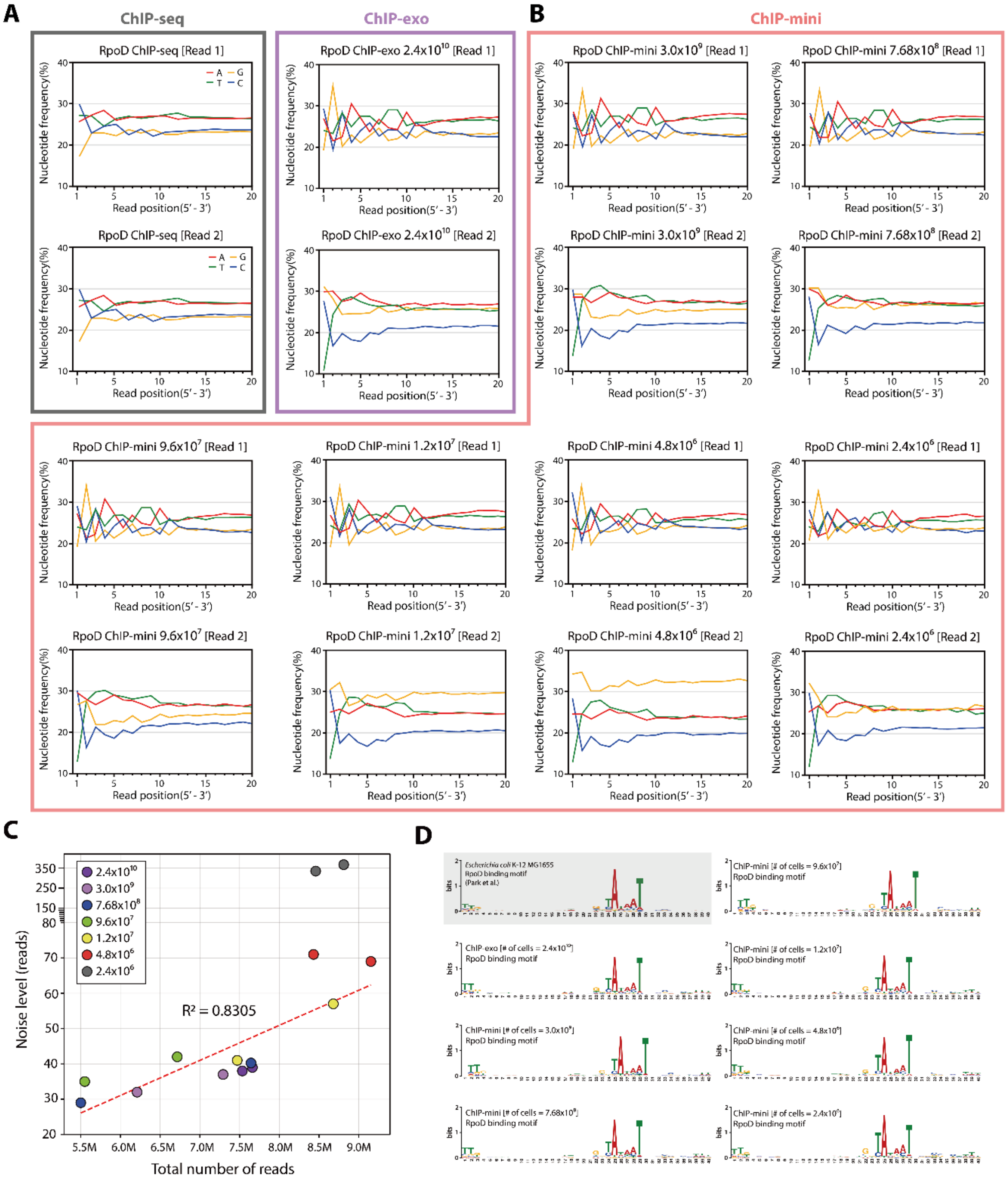


**Figure S2. Comparison of *E. coli* RpoD ChIP-mini datasets according to different number of initial cells.** (A) Nucleotide frequency at the 5' end of the paired-end sequencing reads for ChIP-seq and ChIP-exo libraries. In the ChIP-seq library, read_1 and read_2 exhibit adapter ligation after dA-tailing of sonicated ends, while ChIP-exo library show the product of the exonuclease-digested 5' end in Read_1 file. (B) Nucleotide frequency at the 5' end of the paired-end sequencing reads for ChIP-mini libraries. Read_1 files of ChIP-mini libraries show the product of the exonuclease-digested 5' end, which is a trait of a ChIP-exo library. (C) Determination of noise level in ChIP-exo and ChIP-mini sequencing reads. Noise level was correlated with the total number of reads until reaching $4.8x10^6$ initial bacterial cells. (D) Motif analysis of RpoD binding sites was performed on traditional ChIP-exo and each ChIP-mini dataset, resulting in identical sequence motifs.

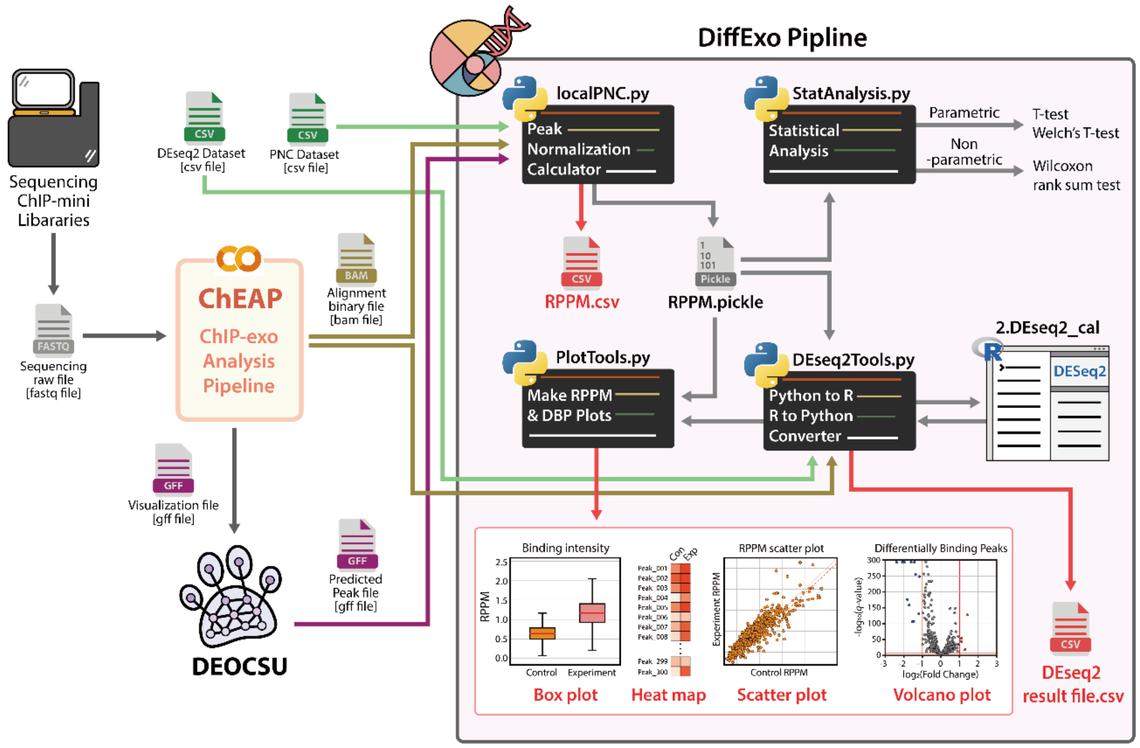


**Figure S3. Schematic of the DiffExo pipeline.** Two dataset files (CSV), including information on the data directories and statistical groups (control or experiment), are formatted before starting the DiffExo pipeline. For preprocessing of ChIP-mini data for the DiffExo pipeline, the sequencing raw files (FASTQ) are converted into alignment files (BAM) and visualization files (GFF) using the cloud-based ChIP-exo analysis pipeline, ChEAP [8]. Followed by visualization, a Deep-learning optimized ChIP-exo peak calling suite (DEOCSU) is utilized to predict binding peaks of target DNA-binding protein using GFF files [9]. Since all the read count data from each ChIP-mini sequencing library are not normally distributed as they are in an RNA-seq library, a negative binomial distribution-based algorithm was adopted to estimate differentially binding sites [10]. In the DiffExo pipeline, 1) localPNC.py: calculate the normalized binding intensity of DNA-binding protein using an RPPM unit and generate a pickle file to convey the normalized data to the next process. 2) StatAnalysis.py: proceed statistical test (parametric or non-parametric test) using the RPPM.pickle file. 3) DEseq2Tools.py: convert overlapping binding sites into reference file for DEseq2, and differential binding sites are calculated in R script (DEseq2_cal). In addition, the raw result file from DEseq2 merges with RPPM and information on binding sites to generate the final Deseq2 result file (CSV) using DEseq2Tools. 4) PlotTools.py: generate four types of plots: box plots, heat maps, scatter plots for visualizing binding intensity, and volcano plots for visualizing DEseq2 results.

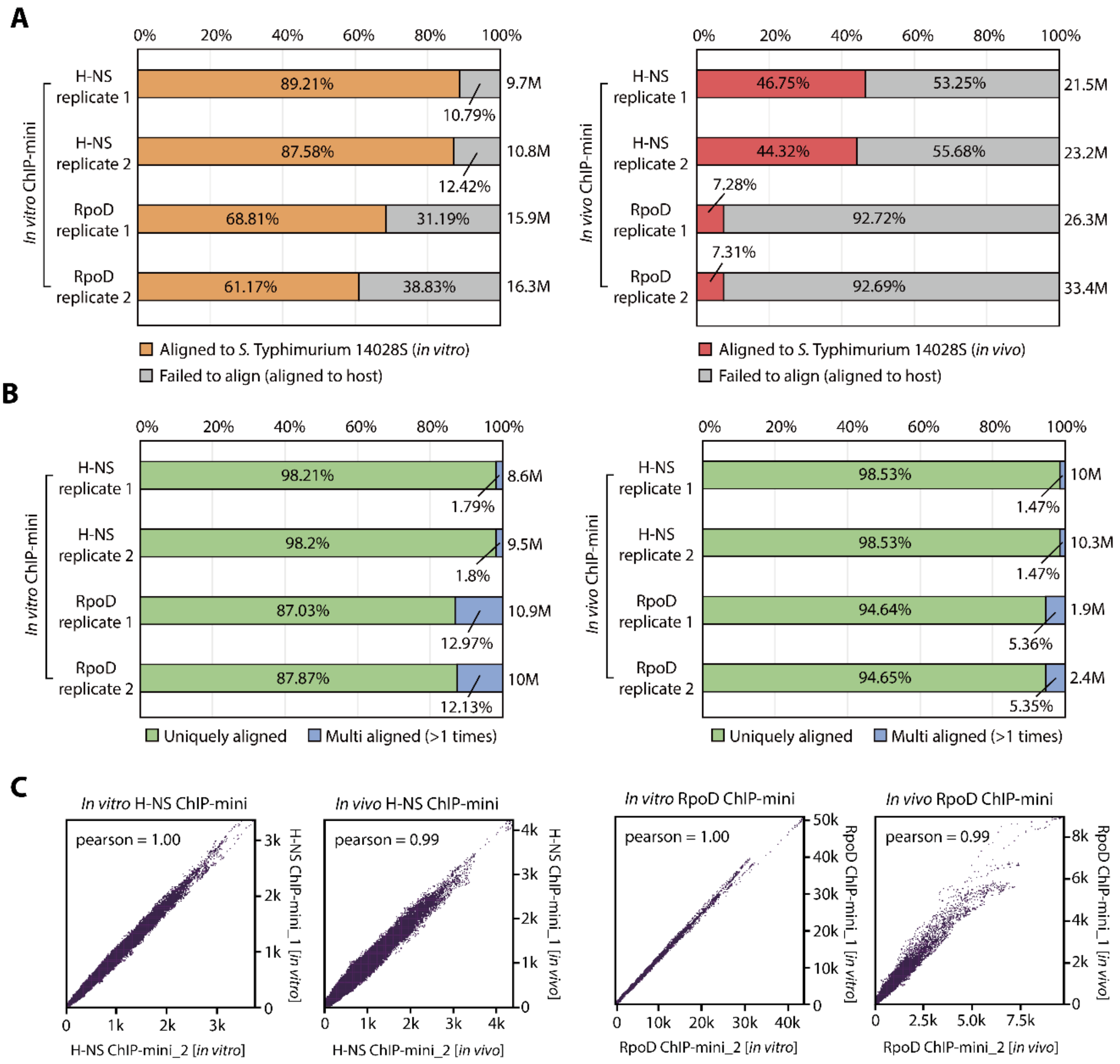


**Figure S4. Detail analysis of sequencing libraries of H-NS and RpoD using *in vitro* and *in vivo* ChIP-mini methods.** (A) Percentage of aligned and failed-aligned reads on the genome of *S.* Typhimurium in H-NS and RpoD libraries. Failed-aligned reads were confirmed to align the host genome (*Mus* musculus). (B) Percentage of uniquely mapping reads and multi aligned reads on the genome of *S.* Typhimurium in H-NS and RpoD libraries. (C) Correlation between replicates of ChIP-mini libraries for H-NS and RpoD was measured by Pearson coefficient. Read counts of each library were split into 10 bp-bins across the *S.* Typhimurium genome, and each dot in the scatter plots represents one genome region.

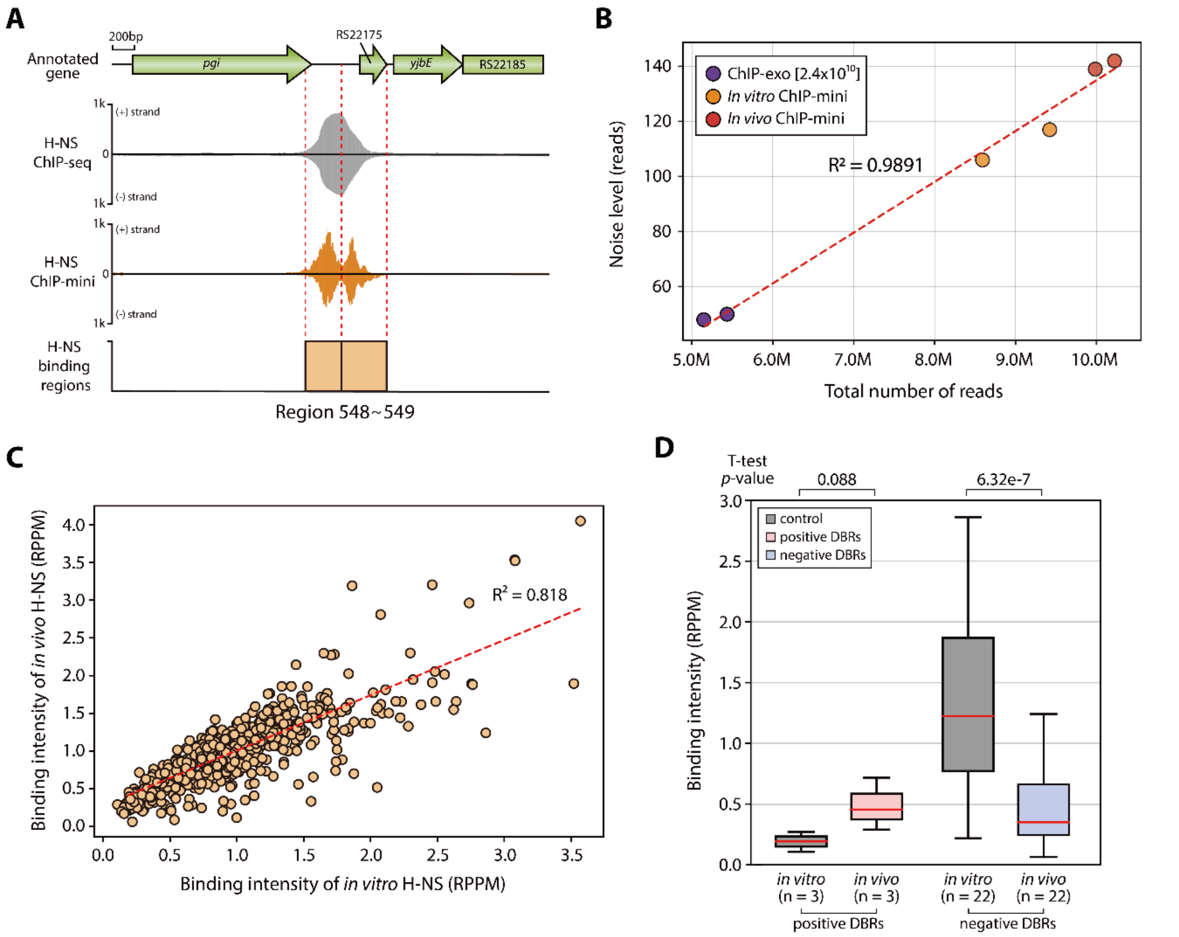


**Figure S5. Detail analysis of H-NS binding regions using ChIP-mini applications.** (A) Comparison of H-NS binding regions from ChIP-seq and ChIP-mini. ChIP-mini also provides better resolution compared to ChIP-seq to dissect binding region, even for the H-NS dataset. ChIP-seq and ChIP-mini for H-NS of *S.* Typhimurium were conducted under M9 minimal media. (B) Determination of noise level in H-NS ChIP-exo and ChIP-mini sequencing reads. (C) Pearson correlation coefficient of normalized H-NS binding intensities was calculated from the *in vitro* and *in vivo* ChIP-mini datasets. (D) A boxplot of DBR binding intensity from *in vitro* and *in vivo* ChIP-mini datasets. The *in vitro* ChIP-mini dataset was used as a control.

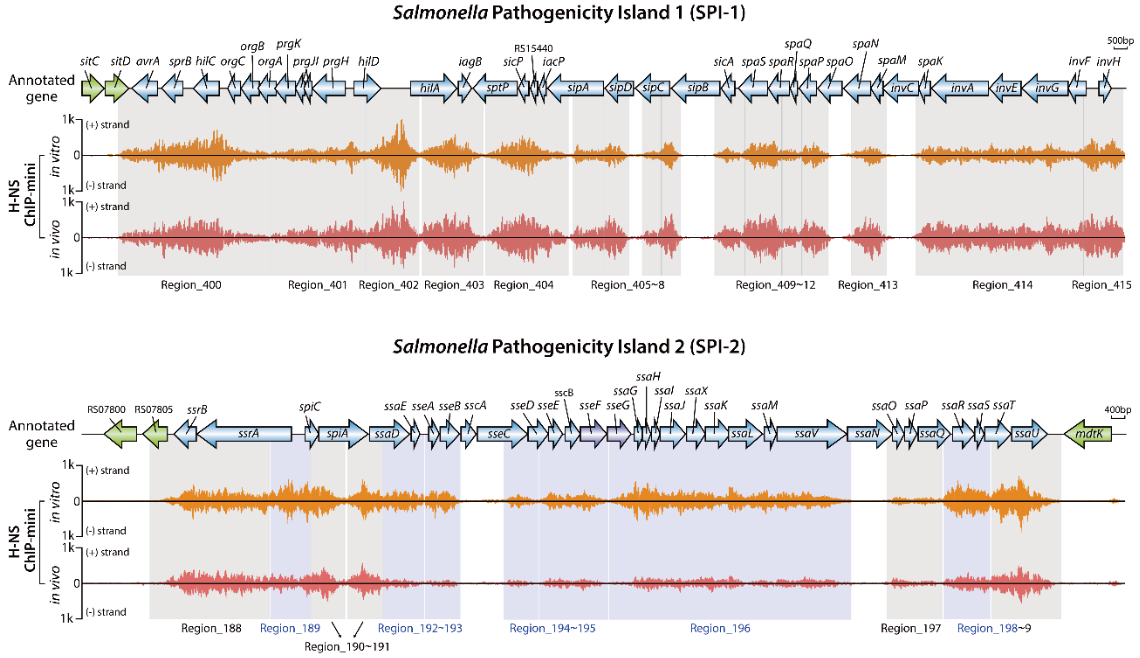


**Figure S6. Overviews of H-NS binding profiles in SPI-1 and SPI-2 of *S.* Typhimurium within macrophages.** Blue boxes denote negative DBRs, while grey boxes indicate non-DBRs. (+) and (−) strands in ChIP-mini data indicate reads mapped on forward and reverse strands, respectively.

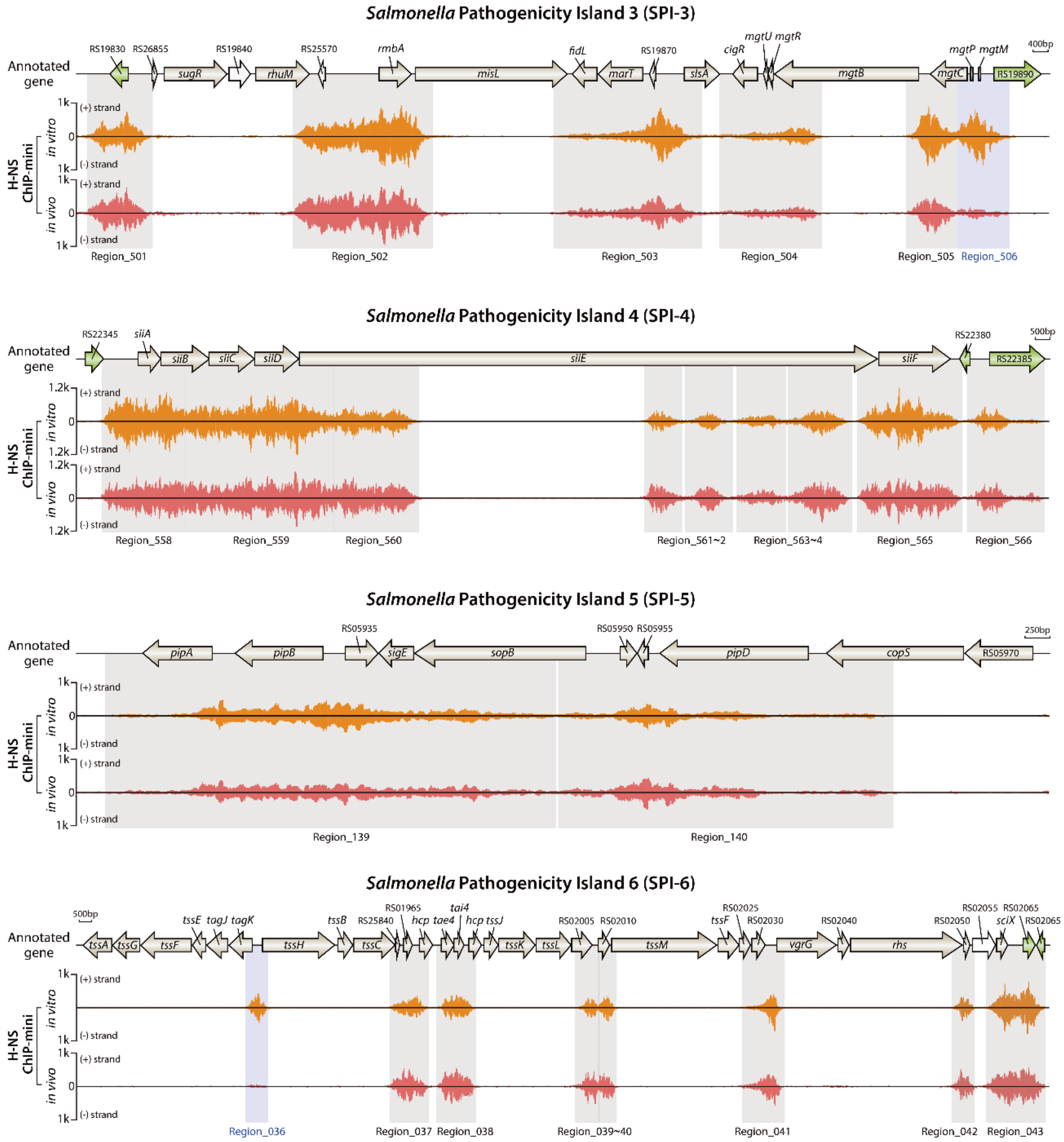


**Figure S7. Overviews of H-NS binding profiles in SPI-3~6 of *S.* Typhimurium within macrophages.** Blue boxes denote negative DBRs, while grey boxes indicate non-DBRs. (+) and (−) strands in ChIP-mini data indicate reads mapped on forward and reverse strands, respectively.

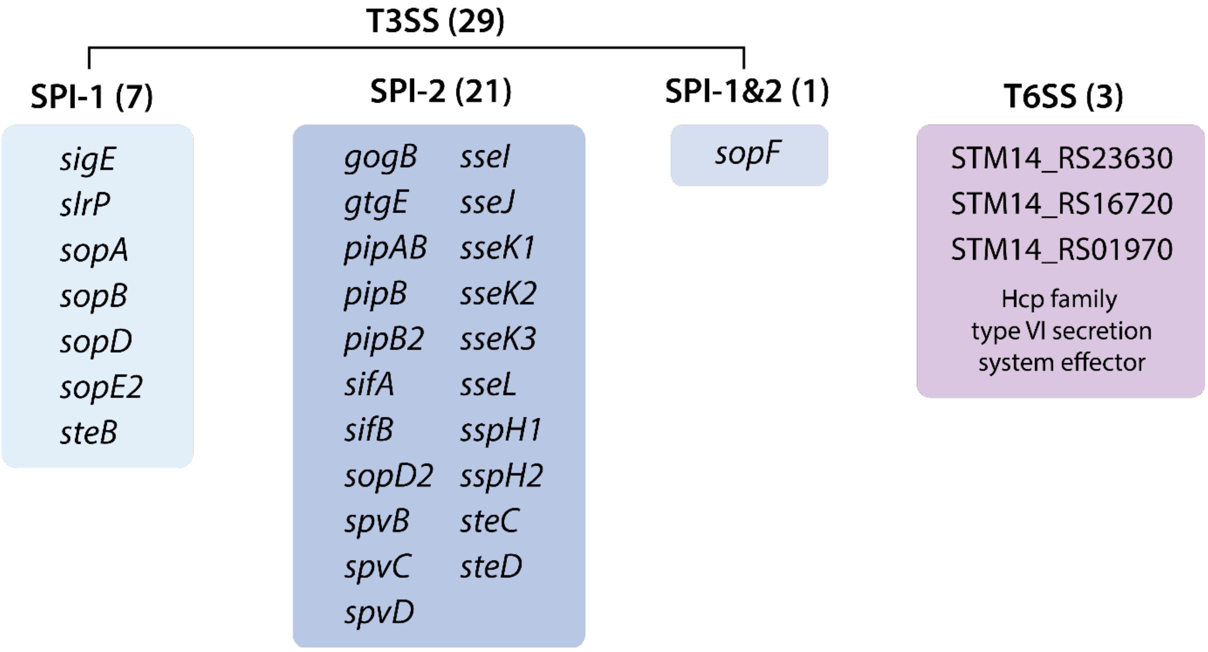


**Figure S8. T3SS and T6SS effector genes associated with H-NS binding regions.** Seven effector genes are related to the SPI-1 T3SS, and 21 effector genes are related to the SPI-2 T3SS. Additionally, *sopF* is associated with both T3SS systems. Three effector genes are associated with the T6SS, encoding the Hcp family effector.

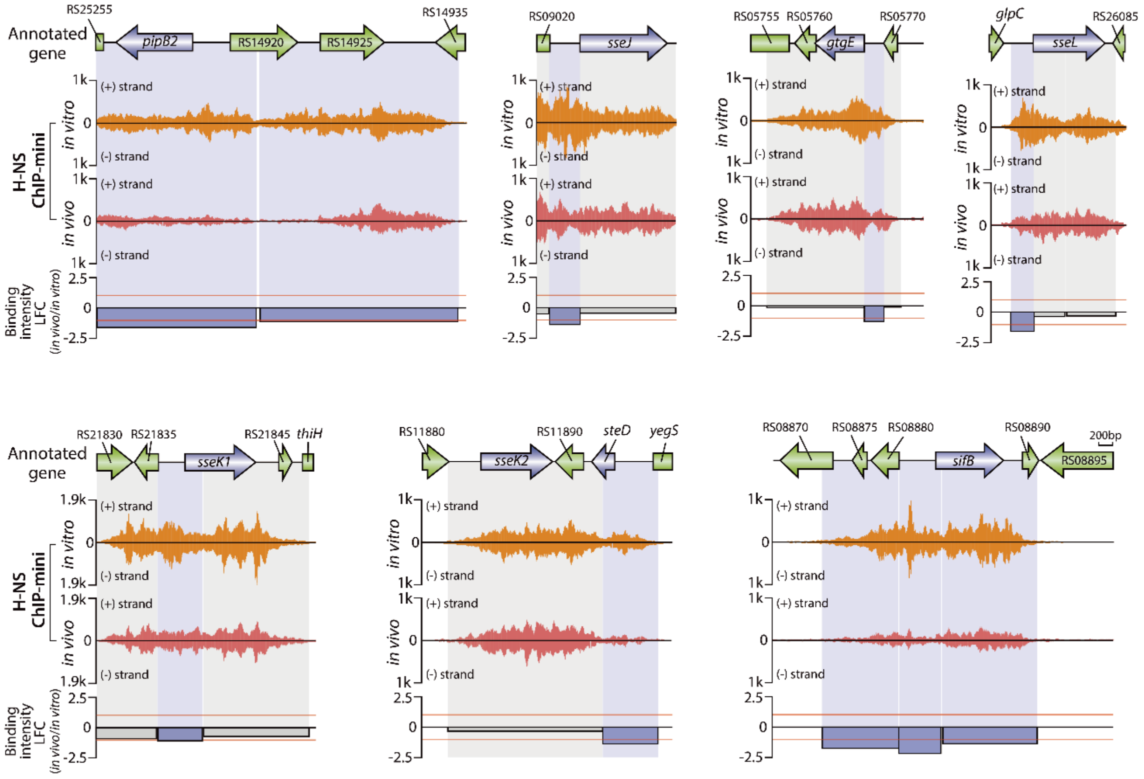


**Figure S9. Negative H-NS DBRs on seven SPI-2 effector genes of *S.* Typhimurium.** Under macrophage intracellular conditions, H-NS binding intensity showed a more significant reduction in the intergenic regions of SPI-2 effector genes. The red lines in binding intensity LFC represent the threshold for DBRs, indicated by -1 and 1. Blue boxes denote negative DBRs, while grey boxes indicate non-DBRs. (+) and (−) strands in ChIP-mini data indicate reads mapped on forward and reverse strands, respectively.

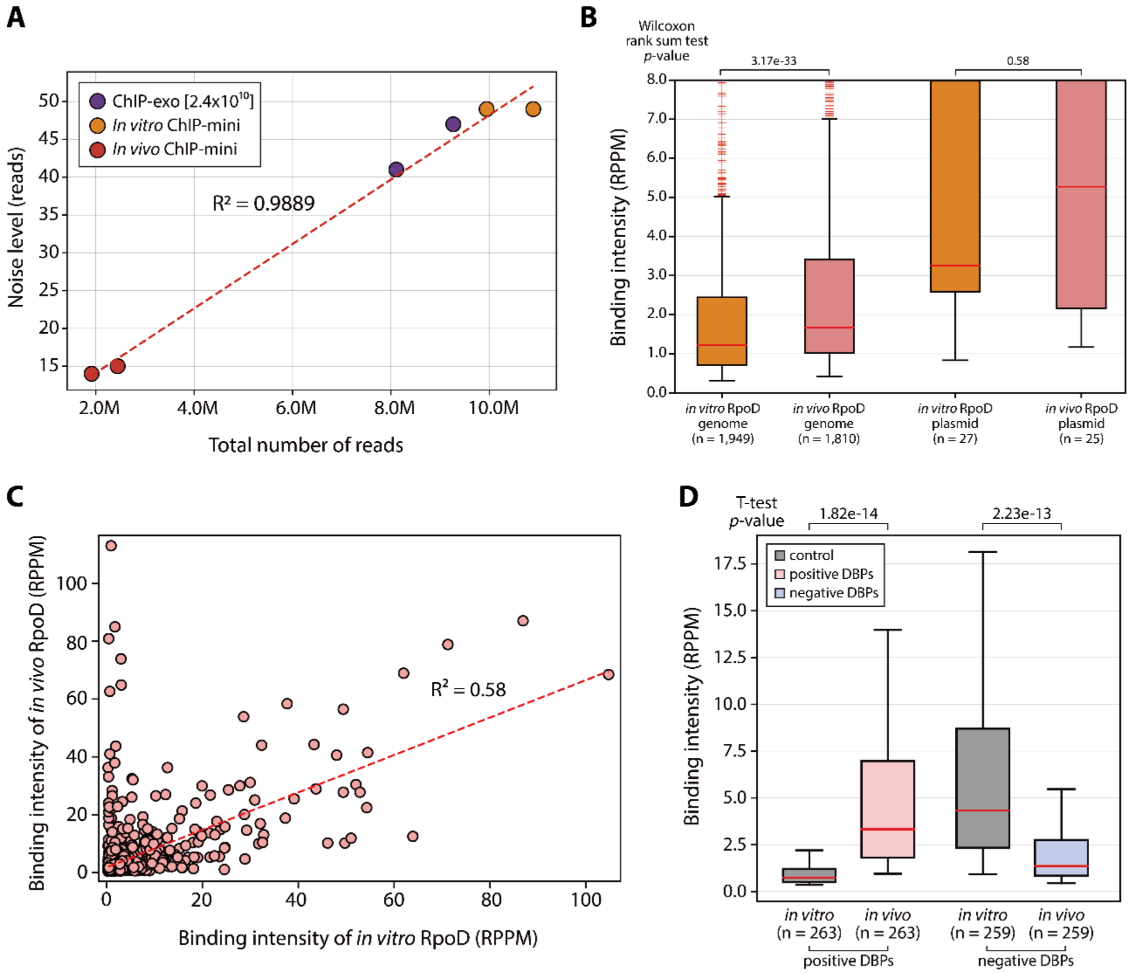


**Figure S10. Detail analysis of RpoD binding sites using ChIP-mini applications.** (A) Determination of noise level in RpoD ChIP-exo and ChIP-mini sequencing reads. (B) Normalized binding intensities of total binding sites were calculated using *in vitro* and *in vivo* RpoD ChIP-mini datasets. (C) Pearson correlation coefficients of normalized RpoD binding intensities were calculated from the overlapping binding sites. (F) Boxplot of DBP binding intensity from *in vitro* and *in vivo* ChIP-mini datasets. The *in vitro* ChIP-mini dataset was used as a control.

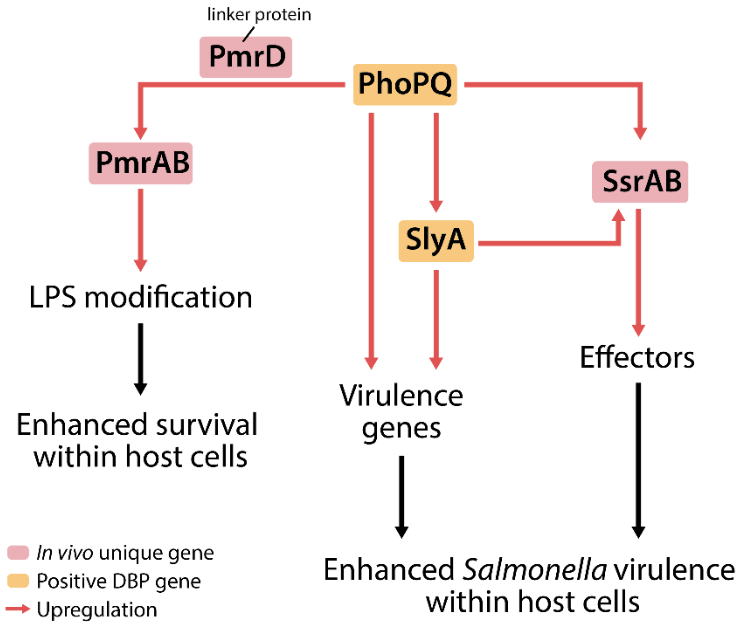


**Figure S11. Regulatory cascades activating the expression of SPI-2 effectors and LPS modification genes.** Pink boxes denote *in vivo* unique RpoD binding gene, and orange box indicates positive DBP gene.

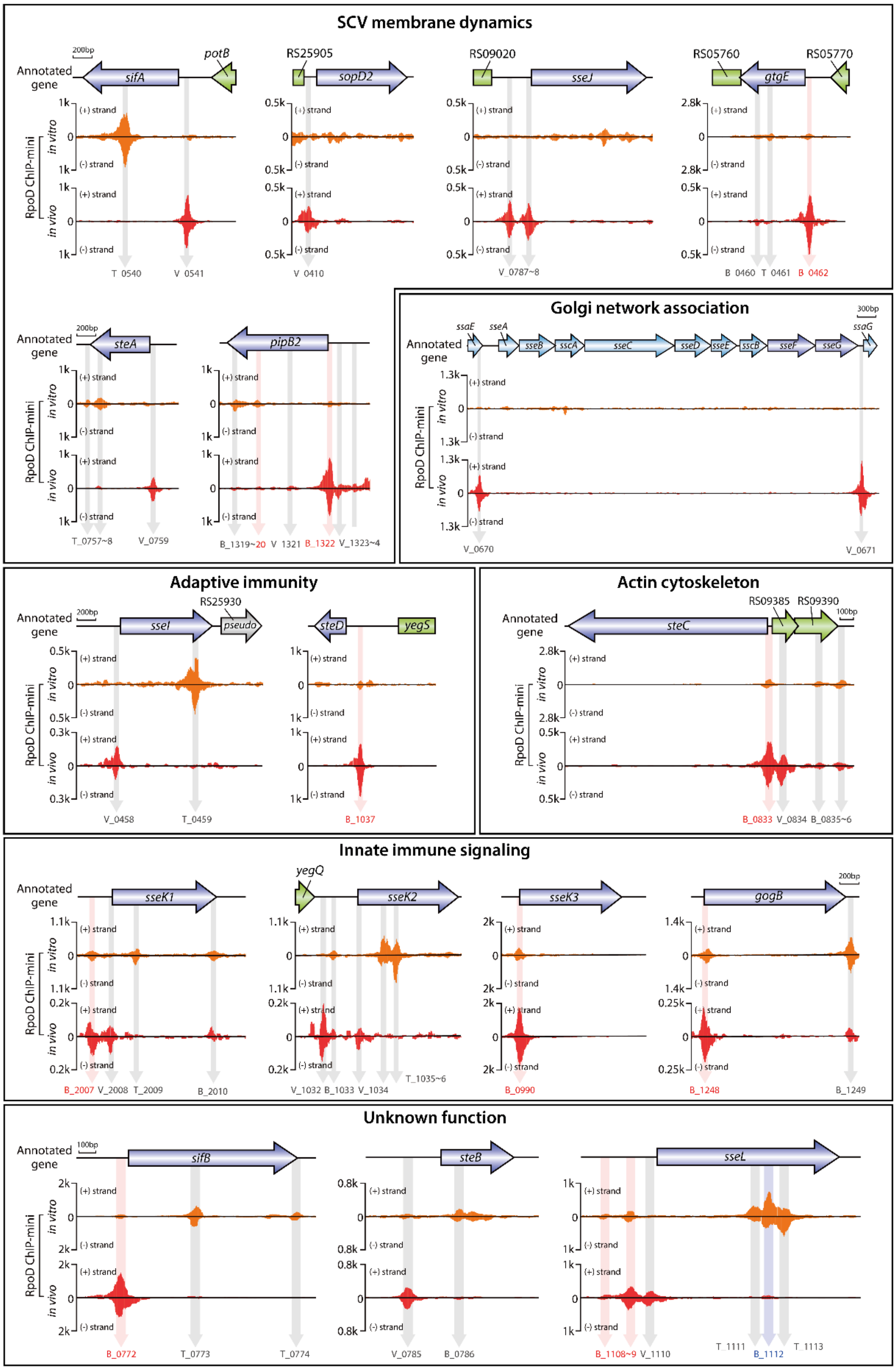


**Figure S12. Overview of *S.* Typhimurium SPI-2 effector genes with RpoD binding profiles under macrophage intracellular conditions.** SPI-2 effector genes were classified based on Jennings et al [11].

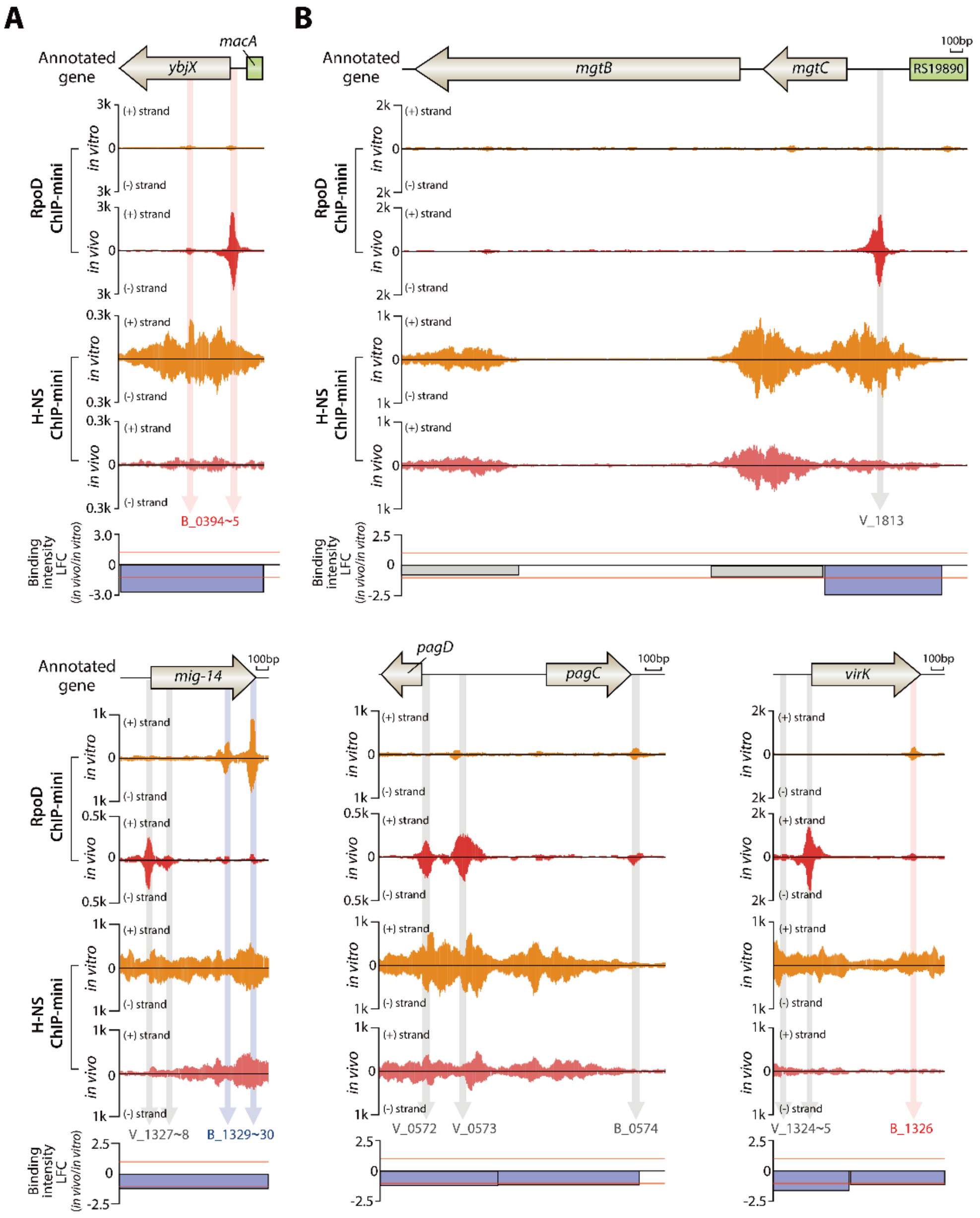


**Figure S13. Dynamic changes in RpoD binding sites on other virulence genes under the influence of H-NS negative DBRs.** (A) *ybjX* was identified as overlapping virulence genes between RpoD positive DBP genes and H-NS negative DBR genes. (B) Five overlapping virulence genes between *in vivo* unique RpoD genes and H-NS negative DBR genes.

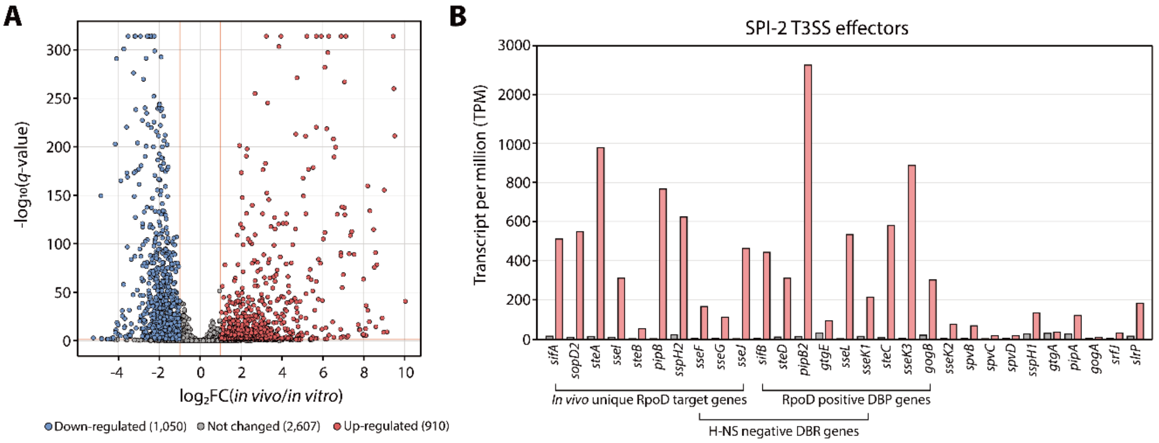


**Figure S14. Comparison of transcript expression levels between macrophage extracellular and intracellular conditions.** (A) Volcano plot displaying differentially expressed genes (DEGs) as response to environmental changes from macrophage extracellular and intracellular conditions (absolute $\log_2$ fold change $\geq 1$, and false discovery rate $< 0.05$). (B) The mRNA expression level of SPI-2 effector genes under both conditions.

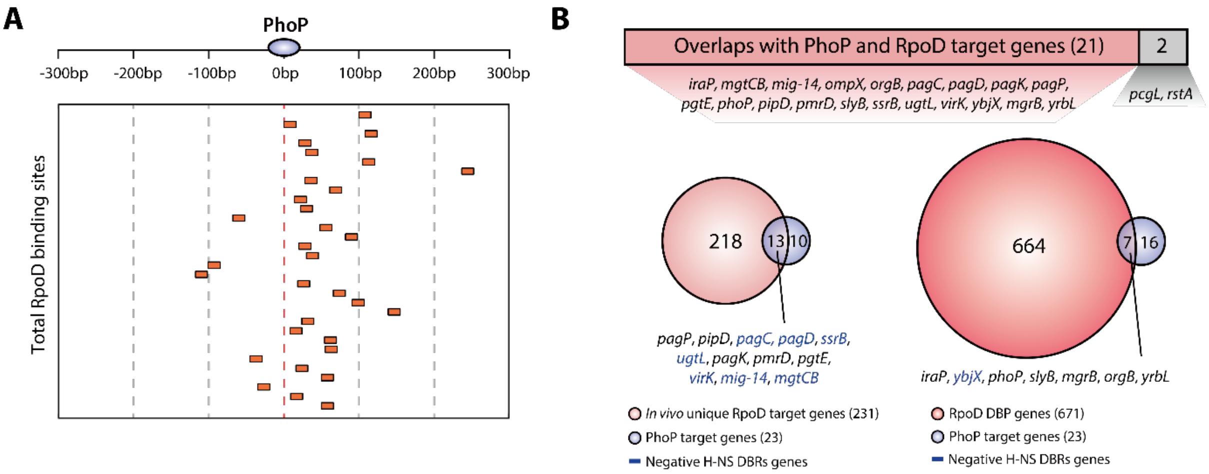


**Figure S15. Comparison of PhoP binding sites with RpoD and H-NS binding sites.** (A) The majority of PhoP binding sites were found in upstream of RpoD. (B) Analysis of the overlapping target genes between PhoP and RpoD under macropahge intracelluakr conditions. 21 out of 23 PhoP target genes were identified among *in vivo* unique RpoD target genes and DBP genes. The gene *ompX* was found in non-DBP genes under both conditions. Genes denoted in blue font are associated with H-NS negative DBRs.

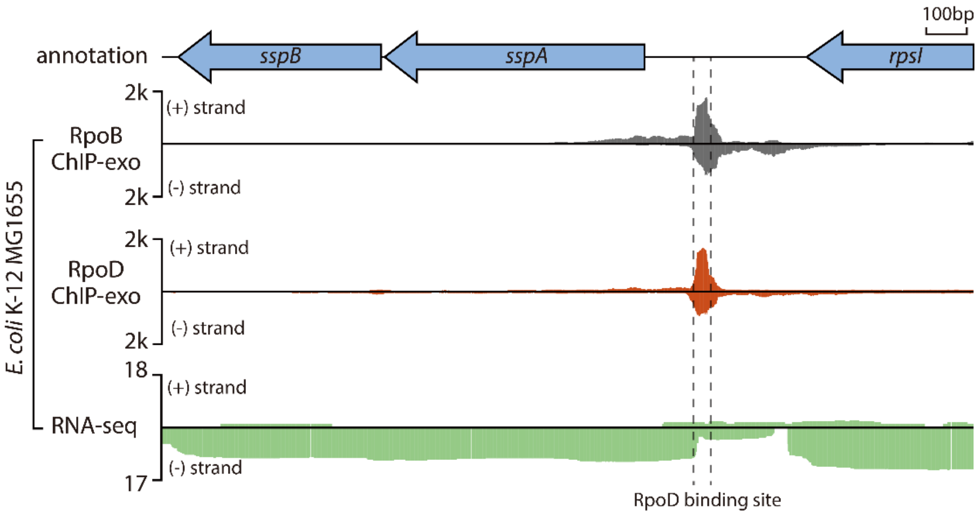


**Figure S16. ChIP-exo bindings of RpoD and RpoB upstream of *sspAB* in *E. coli*.**

**Supplementary Methods**

▶ **ChIP-mini (ChIP-exo minimization for bacterial cells)**

**1. Crosslinking Preparation**

- Prepare crosslinking mix as below:

| Components | 6.25 ml | 1.6 ml | 200 µl |
|---|---|---|---|
| Formaldehyde (37% w/w) | 175 µl | 44 µl | 6 µl |
| TBS (pH 7.4) | 6.5 ml | 1.6 ml | 200 µl |

*Incubate for 25 min at RT, spin down and wash 3 time with ice-cold TBS.*

For samples with a culture volume of less than 200 µl (25 µl, 10 µl, and 5 µl), crosslinking cells were prepared through serial dilution, starting with a 200 µl sample.

**2. Fragmentation and Preparation of Antibody-TF complex**

- Prepare lysis buffer as below:

| Components | Volume |
|---|---|
| 1 M Tris-HCl (pH 7.5) | 5 mL |
| 5 M NaCl | 10 mL |
| 0.5 M EDTA | 1 mL |
| Nuclease-free TDW | 484 mL |
| **Total Volume** | **500 mL** |

*Note: The final concentration of lysis buffer is 10 mM Tris-HCl (pH 7.5), 100 mM NaCl and 1 mM EDTA.*

- Prepare protease inhibitor cocktail (PIC) mix as blow:

| Components | Volume |
|---|---|
| 1 M Tris-HCl (pH 7.5) | 50 mg |
| DMSO | 250 µl |
| Nuclease-free TDW | 750 µl |
| **Total Volume** | **1000** µl |

- Prepare IP buffer as below:

| Components | Volume |
|---|---|
| 1 M Tris-HCl (pH 7.5) | 50 mL |
| 5 M NaCl | 20 mL |
| 0.5 M EDTA | 1 mL |
| Triton X-100 | 10 mL |
| Nuclease-free TDW | 419 mL |
| **Total Volume** | **500 mL** |

*Note: The final concentration of IP buffer is 100 mM Tris-HCl (pH 7.5), 200 mM NaCl, 2% Triton X-100 and 1 mM EDTA.*

- Prepare **washing buffers** as below:

① Washing buffer 1: 50 mM Tris-HCl (pH 7.5), 140 mM NaCl, 1% Triton X-100 and 1mM EDTA

② Washing buffer 2: 50 mM Tris-HCl (pH 7.5), 500 mM NaCl, 1% Triton X-100 and 1mM EDTA

③ Washing buffer 3: 10 mM Tris-HCl (pH 8.0), 250 mM LiCl, 1% Triton X-100 and 1mM EDTA

④ Washing buffer 4 (TE buffer): 10 mM Tris-HCl (pH 8.0), 1mM EDTA

- Prepare fragmentation buffer as blow:

| Components | 6.25 ml | 1.6 ml | 200 ~ 5µl |
|---|---|---|---|
| lysis buffer | 125 µl | | 100 µl |
| PIC mix | 10 µl | | 8 µl |
| Lysozyme (10 mg/ml) | 0.25 µl | | 0.2 µl |
| IP buffer | 137.5 µl | | 110 µl |
| Wash buffer 1 | 75 µl | | 60 µl |

2-3) Resuspend the cell pellet in lysis buffer.

2-4) Add PIC mix and lysozyme.

2-5) Incubate for 30 min at 37 °C on a rotator.

2-6) Add IP buffer and incubate 30 min at 4 °C.

2-7) Shear the lysate by sonication for 30~40 minutes, amplitude 50%, 50” on/10” off, cooling at 4 °C.

- Add 1st antibody to the chromatin solution as blow:

| Components | 6.25 ml | 1.6 ml | 200 ~ 5µl |
|---|---|---|---|
| 1st Antibody (RpoD antibody) | 3 µl | | 1.5 µl |
| 1st Antibody (c-Myc antibody) | 7.5 µl | | 3.75 µl |

*Continue to incubate 6~8 hours at 4 °C with rotating.*

**3. Antibody-TF complex Binding to Dynabeads**

**-** Prepare elution buffer as below:

| Components | Volume |
|---|---|
| 1 M Tris-HCl (pH 7.5) | 25 ml |
| SDS | 5 g |
| 0.5 M EDTA | 1 ml |
| Nuclease-free TDW | 474 ml |
| **Total Volume** | **500 ml** |

*Note: The final concentration of elution buffer is 50 mM Tris-HCl (pH 8.0), 1% SDS and 1 mM EDTA.*

- Prepare bead washing solution as blow:

| Components | Volume |
|---|---|
| BSA powder | 250 mg |
| Ice-cold PBS | 50 ml |
| **Total Volume** | **50 ml** |

- Prepare Dynabeads Pan mouse IgG as blow:

| | 6.25 ml | 1.6 ml | 200 ~ 5μl |
|---|---|---|---|
| Dynabeads Pan mouse IgG | 30 μl | | 15 μl |

*Note: Pull down the beads with the MPC magnet for 30 seconds after washing.*

3-1) Wash with 0.5 ml of bead washing solution 3 times.

3-2) Add Dynabeads Pan mouse IgG to the chromatin solution and incubate overnight at 4 °C with rotating.

**4. STAGE 1: Exonuclease Digestion of Antibody-TF complex**

- Wash the beads with wash buffers as blow:

| | 6.25 ml | 1.6 ml | 200 ~ 5μl |
|---|---|---|---|
| Wash buffer 1 (2 times) | | 500 μl | |
| Wash buffer 2 | | 500 μl | |
| Wash buffer 3 | | 500 μl | |
| Wash buffer 4 | | 500 μl | |

1) End Repair

| Components | 6.25 ml | 1.6 ml | 200 ~ 5μl |
|---|---|---|---|
| End Repair Buffer (10X) | 2.5 μl | | 1 μl |
| End Repair Enzyme Mix | 1.25 μl | | 0.5 μl |
| Water | 21.25 μl | | 8.5 μl |
| **Total Volume** | **25 μl** | | **10 μl** |

*Incubate in a thermal cycler for 30 min at 20 °C and wash the beads with wash buffers.*

2) dA-Tailing

| Components | 6.25 ml | 1.6 ml | 200 ~ 5μl |
|---|---|---|---|
| Water | 10.5 μl | | 4.2 μl |
| dA-Tailing Buffer (10X) | 1.25 μl | | 0.5 μl |
| Klenow Fragment (exo$^-$) | 0.75 μl | | 0.3 μl |
| **Total Volume** | **12.5 μl** | | **5 μl** |

*Incubate in a thermal cycler for 30 min at 37 °C and wash the beads with wash buffers.*

3) Ligation of First Adapter

| Components | 6.25 ml | 1.6 ml | 200 ~ 5µl |
|---|---|---|---|
| Quick Ligation Buffer (2X) | 6.25 µl | | 2.5 µl |
| First Adapter (15 µM) | 1.25 µl | | 0.5 µl |
| Quick T4 DNA Ligase | 0.25 µl | | 0.1 µl |
| Water | 4.75 µl | | 1.9 µl |
| **Total Volume** | **12.5 µl** | | **5 µl** |

*Incubate in a thermal cycler for 15 min at 20 °C and wash the beads with wash buffers.*

4) Nick Repair with NEB PreCR Repair Mix

| Components | 6.25 ml | 1.6 ml | 200 ~ 5µl |
|---|---|---|---|
| Water | 10.75 µl | | 4.3 µl |
| ThermoPol Buffer (10X) | 1.25 µl | | 0.5 µl |
| 10 mM dNTPs | 0.125 µl | | 0.05 µl |
| $NAD^+$ (100X) | 0.125 µl | | 0.05 µl |
| PreCR Mix | 0.25 µl | | 0.1 µl |
| **Total Volume** | **12.5 µl** | | **5 µl** |

*Incubate the repair reaction at 37 °C for 15-20 min and wash the beads with wash buffers.*

5) λ Exonuclease Treatment

| Components | 6.25 ml | 1.6 ml | 200 ~ 5µl |
|---|---|---|---|
| Water | 10.75 µl | | 4.3 µl |
| λ Exonuclease Reaction Buffer (10X) | 1.25 µl | | 0.5 µl |
| λ Exonuclease (5,000 U/mL) | 0.5 µl | | 0.2 µl |
| **Total Volume** | **12.5 µl** | | **5 µl** |

*Incubate at 37 °C for 30 min and wash the beads with wash buffers.*

6) RecJ Exonuclease Treatment

| Components | 6.25 ml | 1.6 ml | 200 ~ 5µl |
|---|---|---|---|
| Water | 11 µl | | 4.4 µl |
| NEBuffer 2 (10X) | 1.25 µl | | 0.5 µl |
| RecJ Exonuclease (30,000 U/mL) | 0.25 µl | | 0.1 µl |
| **Total Volume** | **12.5 µl** | | **5 µl** |

*Incubate at 37 °C for 30 min and wash the beads with wash buffers.*

7) Elution of Dynabeads

| Components | 6.25 ml | 1.6 ml | 200 ~ 5µl |
|---|---|---|---|
| Elution buffer | 50 µl | | 20 µl |

*Continue to incubate overnight at 65 °C.*

**5. Reverse Crosslinking and DNA Purification**

- Pull down the beads with the MPC magnet for 1 min and save the supernatant.

1) RNA Removal
- Prepare RNA removal solution as blow:

| **Components** | **6.25 ml** | **1.6 ml** | **200 ~ 5μl** |
|---|---|---|---|
| RNaseA solution (100 mg/ml RNaseA) | 0.25 μl | | 0.1 μl |
| Washing buffer 4 | 0.75 μl | | 0.9 μl |
| **Total Volume** | **1 μl** | | **1 μl** |

*Add 1 μl of RNA removal solution and incubate at 37 °C for 2 hours.*

2) Reverse Crosslinking
- Prepare protein removal solution as blow:

| **Components** | **6.25 ml** | **1.6 ml** | **200 ~ 5μl** |
|---|---|---|---|
| Protease K (20 mg/ml) | 1 μl | | 0.4 μl |
| Washing buffer 4 | 3 μl | | 3.6 μl |
| **Total Volume** | **4 μl** | | **4 μl** |

*Add 4 μl of protein removal solution and incubate at 55 °C for 2 hours.*

3) IP-DNA Purification
- Add 2.5X DNA purification beads (AccuBead or AMPure beads) to the IP-DNA solution as blow:

| **Components** | **6.25 ml** | **1.6 ml** | **200 ~ 5μl** |
|---|---|---|---|
| IP-DNA solution | 55 μl | | 22 μl |
| DNA purification beads | 137.5 μl | | 55 μl |
| **Total Volume** | **192.5 μl** | | **77 μl** |

*Note: Pull down the beads with the MPC magnet for 2 min.*

3-1) Incubate at RT for 10 min and wash the beads with 200 μl of 80% ethanol twice.

3-2) Ensure dried-beads are completely rehydrated and resuspended using 12 μl of nuclease-free water and collect 11 μl of exonuclease treated IP-DNA.

**6. STAGE 2: Construction of Sequencing Library**

1) Second Strand Synthesis Using Specific Primer and dNTPs

- Prepare the following reaction mix:

| **Components** | **Volume** |
|---|---|
| Exonuclease Treated IP DNA | **11 μl** |
| phi29 DNA Polymerase Buffer (10X) | 2 μl |
| BSA (1 μg/ml) | 4 μl |
| dNTPs (10 mM) | 1 μl |
| Second Strand Synthesis Primer (20 μM) | 1 μl |
| **Total Volume** | **19 μl** |

1-1) Incubate in a thermal cycler for 10 min at 95 °C followed by 5 min at 58 °C.

1-2) Allow to cool to RT by 2 min (primer annealing).

1-3) Add 1 µl phi29 DNA polymerase (10 U/ µl) and incubate for 20 min at 30 °C (primer extension) followed by 10 min at 65 °C (heat inactivation).

2) dA-Tailing

2-1) Add 2.5X DNA purification beads (50 µl) in PCR product (20 µl).

2-2) Incubate at RT for 10 min and wash the beads with 200 µl of 80% ethanol twice.

2-3) Ensure dried-beads are completely rehydrated and resuspended using 10 µl of dA-tailing buffer as blow:

| Components | Volume |
|---|---|
| Water | 8.4 µl |
| dA-Tailing Buffer (10X) | 1 µl |
| Klenow Fragment (exo$^-$) | 0.6 µl |
| **Total Volume** | **10 µl** |

*Incubate at 37 °C for 30 min followed by 30 min enzyme inactivation.*

3) Ligation of Second Adapter

- Prepare the following reaction mix:

| Components | Volume |
|---|---|
| Beads with dA-tailed IP- DNA | 10 µl |
| Quick ligation buffer (2X) | 12.5 µl |
| Second Adapter (15 µM) | 0.625 µl |
| Quick T4 DNA Ligase | 0.625 µl |
| Water | 1.25 µl |
| **Total Volume** | **25 µl** |

*Incubate in a thermal cycler for 15 min at 20 °C.*

3-1) Add 1X PEG/NaCl solution (25 µl) in beads with adapter-ligated IP-DNA (25 µl).

3-2) Incubate at RT for 10 min and wash the beads with 200 µl of 80% ethanol twice.

3-3) Ensure dried-beads are completely rehydrated and resuspended using 11 µl of nuclease-free water and collect 10 µl of adapter-ligated IP DNA.

4) Remove 3' overhang with T4 polymerase

- Prepare the following reaction mix:

| Components | Volume |
|---|---|
| Adapter-ligated IP- DNA | 10 µl |
| Water | 7.4 µl |
| NEBuffer 2.1 | 2 µl |
| dNTPs (10mM) | 0.3 µl |
| T4 DNA polymerase (3 U/ µl) | 0.3 µl |
| **Total Volume** | **20 µl** |

*Incubate in a thermal cycler for 20 min at 12 °C*

4-1) Add 1.0X DNA purification beads (20 µl) in IP-DNA with 2 adapter ligated (20 µl).

4-2) Incubate at RT for 10 min and wash the beads with 200 µl of 80% ethanol twice.

4-3) Ensure dried-beads are completely rehydrated and resuspended using 21 µl of nuclease-free water and collect 20 µl for PCR amplification.

5) PCR Enrichment

| Components | Volume |
|---|---|
| IP-DNA with 2 adapter-ligated | 20 µl |
| KAPA Hifi ReadyMix | 25 µl |
| SYBR Green (10X) | 1 µl |
| Non-Indexed Primer (25 µM) | 1 µl |
| Indexed Primer (25 µM) | 1 µl |
| Water | 2 µl |
| **Total Volume** | **50 µl** |

5-1) Amplify using the following PCR protocol using the qPCR thermocycler:

- 98 °C for 2 min
- 18 cycles of the following:
- 98 °C for 15 s
- 65 °C for 30 s
- 72 °C for 30 s
- 72 °C for 10 min
- Hold at 4 °C

5-2) Add 1.0X DNA purification beads (50 µl) in IP-DNA with PCR reaction mix (50 µl).

5-3) Incubate at RT for 10 min and wash the beads with 200 µl of 80% ethanol twice.

5-4) Ensure dried-beads are completely rehydrated and resuspended using 21 µl of nuclease-free water and collect 20 µl.

5-5) Check library concentration and fragment distribution using the Qubit dsDNA HS kit and Agilent High Sensitivity DNA Kit, respectively.

5-5) Store amplified libraries at -15~20 °C before sequencing.

▶ ***In vitro* and *in vivo* ChIP-mini (ChIP-exo minimization for host-infected bacterial cells)**

**1. *Salmonella* Typhimurium Infection and Crosslinking Preparation**

- Prepare crosslinking mix as below:

| Components | *In vivo* and *in vitro* ChIP-mini |
|---|---|
| Formaldehyde (37% w/w) | 280 μl |
| TBS (pH 7.4) | 10 ml |

1-1) Seed macrophage-like cells (J774A.1) in 75T flasks at a plating density of $6x10^6$ per flask under Dulbecco's modified Eagle's medium (DMEM) supplemented with 10% (v/v) fetal bovine serum (FBS) and antibiotic antimycotic at 37 °C with 5% $CO^2$ in a humidified incubator.

1-2) Inoculate *S.* Typhimurium 14028s from glycerol stock to LB broth and incubate at 37 ℃ overnight with constant agitation.

1-3) Dilute overnight culture into fresh LB broth and culture at 37˚C overnight with agitation in a shaking incubator before infection.

1-4) Add overnight-grown bacteria to the macrophages (macrophage cell number ≈ 8 x $10^6$) at a multiplicity of infection (MOI) of 10, and centrifuge at 500 xg for 5 minutes at RT and incubate for an additional 30 min.

1-5) Transfer 10ml of DMEM including extracellular bacteria in 75T flask and crosslink adding formaldehyde at RT for 25 min (*in vitro* Chip-mini).

1-5-1) Centrifuge crosslinked cells at 4,000 xg and wash 3 time with ice-cold TBS.

1-6) Wash infected-macrophage cells in 75T flask three times with PBS.

1-7) Add 10ml DMEM supplemented with 10% FBS and 150 μg $ml^{-1}$ gentamycin and incubate 37 ℃ for 1 hour.

1-8) Replace 10ml DMEM containing 10% FBS with 15 μg $mL^{-1}$ gentamicin and incubate at 37 ℃ for 6 hours.

1-9) Add formaldehyde to infected-macrophage cells in 75T flask for co-crosslinking at RT for 25 min (*in vivo* ChIP-mini).

1-9-1) Centrifuge at 500 xg co-crosslinked cells and wash 3 time with ice-cold TBS.

1-10) Remove supernatant and transfer co-crosslinked cells to 1.75ml tube.

*Note: The number of intracellular bacteria in the host cells is less than $4.8x10^6$, the efficiency of in vivo ChIP-mini can be significantly reduced.*

**2. Fragmentation and Preparation of Antibody-TF complex**

- Prepare host lysis mix as below:

| Components | *In vivo* ChIP-mini |
|---|---|
| Triton X-100 | 10 μl |
| PBS (pH 7.4) | 990 μl |
| **Total Volume** | **1 ml** |

2-1) Resuspend co-crosslinked cells in 1 ml of host lysis buffer using pipetting and incubate RT at 20 min.

2-2) Centrifuge at 20,000 xg and remove all supernatant in 1.75ml tube (*Do not disturb the cell pellet*).

- Prepare fragmentation buffer as blow:

| Components | *In vitro* and *in vivo* ChIP-mini |
|---|---|
| lysis buffer | 100 µl |
| PIC mix | 8 µl |
| Lysozyme (10 mg/ml) | 0.2 µl |
| IP buffer | 110 µl |
| Wash buffer 1 | 60 µl |

2-3) Resuspend the cell pellet in 100 µl of lysis buffer.

2-4) Add 8 µl of PIC mix and 0.2 µl of lysozyme.

2-5) Incubate for 30 min at 37 °C on a rotator.

2-6) Add 110 µl of IP buffer and incubate 30 min at 4 °C.

2-7) Shear the lysate by sonication for 30~40 minutes, amplitude 50%, 50” on/10” off, cooling at 4 °C.

- Add 1st antibody to the chromatin solution as blow:

| Components | *In vitro* and *in vivo* ChIP-mini |
|---|---|
| 1st Antibody (RpoD antibody) | 1.5 µl |
| 1st Antibody (c-Myc antibody) | 3.75 µl |

*Continue to incubate 6~8 hours at 4 °C with rotating.*

**3. Antibody-TF complex Binding to Dynabeads**

**-** Prepare elution buffer as below:

| Components | Volume |
|---|---|
| 1 M Tris-HCl (pH 7.5) | 25 ml |
| SDS | 5 g |
| 0.5 M EDTA | 1 ml |
| Nuclease-free TDW | 474 ml |
| **Total Volume** | **500 ml** |

*Note: The final concentration of elution buffer is 50 mM Tris-HCl (pH 8.0), 1% SDS and 1 mM EDTA.*

- Prepare bead washing solution as blow:

| Components | Volume |
|---|---|
| BSA powder | 250 mg |
| Ice-cold PBS | 50 ml |
| **Total Volume** | **50 ml** |

- Prepare Dynabeads Pan mouse IgG as blow:

| | *In vitro* and *in vivo* ChIP-mini |
|---|---|
| Dynabeads Pan mouse IgG | 15 µl |

*Note: Pull down the beads with the MPC magnet for 30 seconds after washing.*

3-1) Wash with 0.5 ml of bead washing solution 3 times.

3-2) Add Dynabeads Pan mouse IgG to the chromatin solution and incubate overnight at 4 °C with rotating.

**4. STAGE 1: Exonuclease Digestion of Antibody-TF complex**

- Wash the beads with wash buffers as blow:

| | *In vitro* and *in vivo* ChIP-mini |
|---|---|
| Wash buffer 1 (2 times) | 500 µl |
| Wash buffer 2 | 500 µl |
| Wash buffer 3 | 500 µl |
| Wash buffer 4 | 500 µl |

1) End Repair

| Components | *In vitro* ChIP-mini | *In vivo* ChIP-mini |
|---|---|---|
| End Repair Buffer (10X) | 1 µl | 2 µl |
| End Repair Enzyme Mix | 0.5 µl | 1 µl |
| Water | 8.5 µl | 17 µl |
| **Total Volume** | **10 µl** | **20 µl** |

*Incubate in a thermal cycler for 30 min at 20 °C and wash the beads with wash buffers.*

2) dA-Tailing

| Components | *In vitro* ChIP-mini | *In vivo* ChIP-mini |
|---|---|---|
| Water | 4.2 µl | 8.4 µl |
| dA-Tailing Buffer (10X) | 0.5 µl | 1 µl |
| Klenow Fragment (exo$^-$) | 0.3 µl | 0.6 µl |
| **Total Volume** | **5 µl** | **10 µl** |

*Incubate in a thermal cycler for 30 min at 37 °C and wash the beads with wash buffers.*

3) Ligation of First Adapter

| Components | *In vitro* ChIP-mini | *In vivo* ChIP-mini |
|---|---|---|
| Quick Ligation Buffer (2X) | 2.5 µl | 5 µl |
| First Adapter (15 µM) | 0.5 µl | 1 µl |
| Quick T4 DNA Ligase | 0.1 µl | 0.2 µl |
| Water | 1.9 µl | 3.8 µl |
| **Total Volume** | **5 µl** | **10 µl** |

*Incubate in a thermal cycler for 15 min at 20 °C and wash the beads with wash buffers.*

4) Nick Repair with NEB PreCR Repair Mix

| Components | *In vitro* ChIP-mini | *In vivo* ChIP-mini |
|---|---|---|
| Water | 4.3 µl | 8.6 µl |
| ThermoPol Buffer (10X) | 0.5 µl | 1.0 µl |
| 10 mM dNTPs | 0.05 µl | 0.1 µl |
| $NAD^+$ (100X) | 0.05 µl | 0.1 µl |
| PreCR Mix | 0.1 µl | 0.2 µl |
| **Total Volume** | **5 µl** | **10 µl** |

*Incubate the repair reaction at 37 °C for 15-20 min and wash the beads with wash buffers.*

5) λ Exonuclease Treatment

| Components | *In vitro* ChIP-mini | *In vivo* ChIP-mini |
|---|---|---|
| Water | 4.3 µl | 8.6 µl |
| λ Exonuclease Reaction Buffer (10X) | 0.5 µl | 1 µl |
| λ Exonuclease (5,000 U/mL) | 0.2 µl | 0.4 µl |
| **Total Volume** | **5 µl** | **10 µl** |

*Incubate at 37 °C for 30 min and wash the beads with wash buffers.*

6) RecJ Exonuclease Treatment

| Components | *In vitro* ChIP-mini | *In vivo* ChIP-mini |
|---|---|---|
| Water | 4.4 µl | 8.8 µl |
| NEBuffer 2 (10X) | 0.5 µl | 1 µl |
| RecJ Exonuclease (30,000 U/mL) | 0.1 µl | 0.2 µl |
| **Total Volume** | **5 µl** | **10 µl** |

*Incubate at 37 °C for 30 min and wash the beads with wash buffers.*

7) Elution of Dynabeads

| Components | *In vitro* and *in vivo* ChIP-mini |
|---|---|
| Elution buffer | 20 µl |

*Continue to incubate overnight at 65 °C.*

**5. Reverse Crosslinking and DNA Purification**

- Pull down the beads with the MPC magnet and save supernatant.

1) RNA Removal

- Prepare RNA removal solution as blow:

| Components | *In vitro* and *in vivo* ChIP-mini |
|---|---|
| RNaseA solution (100 mg/ml RNaseA) | 0.1 µl |
| Washing buffer 4 | 0.9 µl |
| **Total Volume** | **1 µl** |

*Add 1 µl of RNA removal solution and incubate at 37 °C for 2 hours.*

2) Reverse Crosslinking
- Prepare protein removal solution as blow:

| Components | *In vitro* and *in vivo* ChIP-mini |
|---|---|
| Protease K (20 mg/ml) | 0.4 µl |
| Washing buffer 4 | 3.6 µl |
| **Total Volume** | **4 µl** |

*Add 4 µl of protein removal solution and incubate at 55 °C for 2 hours.*

3) IP-DNA Purification
- Add 2.5X DNA purification beads (AccuBead or AMPure beads) to the IP-DNA solution as blow:

| Components | *In vitro* and *in vivo* ChIP-mini |
|---|---|
| IP-DNA solution | 22 µl |
| DNA purification beads | 55 µl |
| **Total Volume** | **77 µl** |

*Note: Pull down the beads with the MPC magnet for 2 min.*

3-1) Incubate at RT for 10 min and wash the beads with 200 µl of 80% ethanol twice.

3-2) Ensure dried-beads are completely rehydrated and resuspended using 12 µl of nuclease-free water and collect 11 µl of exonuclease treated IP-DNA.

**6. STAGE 2: Construction of Sequencing Library**

1) Second Strand Synthesis Using Specific Primer and dNTPs

- Prepare the following reaction mix:

| Components | Volume |
|---|---|
| Exonuclease Treated IP DNA | **11 µl** |
| phi29 DNA Polymerase Buffer (10X) | 2 µl |
| BSA (1 µg/ml) | 4 µl |
| dNTPs (10 mM) | 1 µl |
| Second Strand Synthesis Primer (20 µM) | 1 µl |
| **Total Volume** | **19 µl** |

1-1) Incubate in a thermal cycler for 10 min at 95 °C followed by 5 min at 58 °C.

1-2) Allow to cool to RT by 2 min (primer annealing).

1-3) Add 1 µl phi29 DNA polymerase (10 U/ µl) and incubate for 20 min at 30 °C (primer extension) followed by 10 min at 65 °C (heat inactivation).

2) dA-Tailing

2-1) Add 2.5X DNA purification beads (50 µl) in PCR product (20 µl).

2-2) Incubate at RT for 10 min and wash the beads with 200 µl of 80% ethanol twice.

2-3) Ensure dried-beads are completely rehydrated and resuspended using 10 µl of dA-tailing buffer as blow:

| Components | Volume |
|---|---|
| Water | 8.4 µl |
| dA-Tailing Buffer (10X) | 1 µl |
| Klenow Fragment (exo$^-$) | 0.6 µl |
| **Total Volume** | **10 µl** |

*Incubate at 37 °C for 30 min followed by 30 min enzyme inactivation.*

3) Ligation of Second Adapter

- Prepare the following reaction mix:

| Components | Volume |
|---|---|
| Beads with dA-tailed IP- DNA | 10 µl |
| Quick ligation buffer (2X) | 12.5 µl |
| Second Adapter (15 µM) | 0.625 µl |
| Quick T4 DNA Ligase | 0.625 µl |
| Water | 1.25 µl |
| **Total Volume** | **25 µl** |

*Incubate in a thermal cycler for 15 min at 20 °C.*

3-1) Add 1X PEG/NaCl solution (25 µl) in beads with adapter-ligated IP-DNA (25 µl).

3-2) Incubate at RT for 10 min and wash the beads with 200 µl of 80% ethanol twice.

3-3) Ensure dried-beads are completely rehydrated and resuspended using 11 µl of nuclease-free water and collect 10 µl of adapter-ligated IP DNA.

4) Remove 3' overhang with T4 polymerase

- Prepare the following reaction mix:

| Components | Volume |
|---|---|
| Adapter-ligated IP- DNA | 10 µl |
| Water | 7.4 µl |
| NEBuffer 2.1 | 2 µl |
| dNTPs (10mM) | 0.3 µl |
| T4 DNA polymerase (3 U/ µl) | 0.3 µl |
| **Total Volume** | **20 µl** |

*Incubate in a thermal cycler for 20 min at 12 °C.*

4-1) Add 1.0X DNA purification beads (20 µl) in IP-DNA with 2 adapter ligated (20 µl).

4-2) Incubate at RT for 10 min and wash the beads with 200 µl of 80% ethanol twice.

4-3) Ensure dried-beads are completely rehydrated and resuspended using 21 µl of nuclease-free water and collect 20 µl for PCR amplification.

5) PCR Enrichment

| Components | Volume |
|---|---|
| IP-DNA with 2 adapter-ligated | 20 µl |
| KAPA Hifi ReadyMix | 25 µl |
| SYBR Green (10X) | 1 µl |
| Non-Indexed Primer (25 µM) | 1 µl |
| Indexed Primer (25 µM) | 1 µl |
| Water | 2 µl |
| **Total Volume** | **50 µl** |

5-1) Amplify using the following PCR protocol using the qPCR thermocycler:

- 98 °C for 2 min
- 18 cycles of the following:
- 98 °C for 15 s
- 65 °C for 30 s
- 72 °C for 30 s
- 72 °C for 10 min
- Hold at 4 °C

5-2) Add 1.0X DNA purification beads (50 µl) in IP-DNA with PCR reaction mix (50 µl).

5-3) Incubate at RT for 10 min and wash the beads with 200 µl of 80% ethanol twice.

5-4) Ensure dried-beads are completely rehydrated and resuspended using 21 µl of nuclease-free water and collect 20 µl.

5-5) Check library concentration and fragment distribution using the Qubit dsDNA HS kit and Agilent High Sensitivity DNA Kit, respectively.

5-5) Store amplified libraries at -15~20 °C before sequencing.
